# CroCoDeEL: accurate control-free detection of cross-sample contamination in metagenomic data

**DOI:** 10.1101/2025.01.15.633153

**Authors:** Lindsay Goulet, Florian Plaza Oñate, Alexandre Famechon, Benoît Quinquis, Eugeni Belda, Edi Prifti, Emmanuelle Le Chatelier, Guillaume Gautreau

## Abstract

Metagenomic sequencing provides profound insights into microbial communities, but it is often compromised by technical biases, including cross-sample contamination. This phenomenon arises when microbial content is inadvertently exchanged among concurrently processed samples, distorting microbial profiles and compromising the reliability of metagenomic data and downstream analyses. Existing detection methods often rely on negative controls, which are inconvenient and do not detect contamination within real samples. Meanwhile, strain-level bioinformatics approaches fail to distinguish contamination from natural strain sharing and lack sensitivity. To fill this gap, we introduce CroCoDeEL, a decision-support tool for detecting and quantifying cross-sample contamination. Leveraging linear modeling and a pre-trained supervised model, CroCoDeEL identifies specific contamination patterns in species abundance profiles. It requires no negative controls or prior knowledge of sample processing positions, offering improved accuracy and versatility. Benchmarks across three public datasets demonstrate that CroCoDeEL accurately detects contaminated samples and identifies their contamination sources, even at low rates (*<*0.1%), provided sufficient sequencing depth. Notably, we discovered critical contamination cases in highly cited studies, calling some of their results into question. Our findings suggest that cross-sample contamination is a widespread yet underexplored issue in metagenomics and emphasize the necessity of systematically integrating contamination detection into sequencing quality control.

## Introduction

Shotgun metagenomic sequencing has revolutionized microbiology by allowing indepth characterization of microbial communities without prior cultivation in the lab. While being powerful, this technique is subject to technical and experimental biases at different steps, including sample collection and storage, DNA extraction, sequencing and bioinformatics processing [1]. One of these biases is contamination, which refers to the presence in a sample of DNA that does not originate from it. Two types of contamination have been described so far: contaminant DNA and cross-sample contamination [2].

Contaminant DNA can originate from various external sources, including the sampling environment, laboratory settings, DNA extraction kits, or other reagents used during processing. This contamination is often identified by the detection of atypical species that are unexpected in the studied ecosystem. The issue becomes particularly critical when working with low-biomass samples, where contaminant DNA can constitute a substantial proportion of the total DNA in the sample [2, 3].

Cross-sample contamination, also known as well-to-well leakage, refers to the accidental transfer of material between samples processed together. This unintentional mixing of samples performed by a robot or a human operator has several possible causes. These include the use by mistake of the same pipette tip for multiple samples, leakage during pipetting, the formation of micro-droplets when sealing plates with films or even splashing following the agitation of plates or tubes. Cross-sample contamination has been shown to occur primarily between adjacent or physically close samples and is more frequently detected when low and high biomass samples are processed together [4, 5]. Other phenomena occurring during sequencing may cause cross-sample contamination, such as index hopping, referring to the assignment of sequencing reads to the wrong sample in multiplexed libraries [6].

In contrast to contaminant DNA, cross-sample contamination remains underexplored. It is often overlooked by laboratory staff and data analysts, posing a significant barrier to the replicability of studies. If not detected and treated accordingly, this *splashome* [7] can lead to biased results, such as the false discovery of microbial species including pathogens, the misdetection of strain-sharing events [5], potential loss of statistical power, or the overestimation of α-diversity, especially richness [8].

Processing samples in 96-well plates has become a standard practice to maximize throughput. However, this approach drastically increases the risk of cross-sample contamination given the minimal physical separation between wells and the use of sealing films. To mitigate this risk, some laboratory protocols recommend using plates with individual microtubes instead of traditional wells, providing better physical separation between samples [9]. Additional precautions include cooling plates after the thermal lysis step and performing centrifugation to prevent the formation of micro-droplets on sealing films. Moreover, drilling sealing films rather than removing them entirely may further reduce the risk of droplet dispersion (unpublished protocols). Finally, the issue of index hopping can be addressed by employing Unique Dual Index (UDI) sequencing adapters [10].

Wet lab strategies can be employed to detect cross-sample contamination, with negative controls (blanks) being the most common approach. These controls, if uncontaminated, should contain no genetic material [2, 9]. SCRuB [11] leverages sample location data to determine whether negative controls have been contaminated by external sources or well-to-well leakage, enabling accurate removal of external contamination. However, beyond negative controls, SCRuB is not designed to detect cross-sample contamination. While negative controls can reveal contamination during laboratory procedures, they cannot confirm contamination within genuine samples or estimate contamination levels. In addition, contamination events further away from negative controls may remain undetected [5]. Strategically placing multiple negative controls across plates can improve detection [4, 9], but this approach reduces the number of wells available for genuine samples, increasing costs. Many laboratories limit the use of negative controls or fail to publish associated sequencing data, requiring blind trust in public datasets. An alternative strategy is to use spike-ins which are synthetic DNA sequences added to samples. The unexpected detection of spike-in DNA in a sample indicates cross-contamination. However, contamination occurring before addition of spike-ins will remain undetected. Although used in genome sequencing [12], we found no reports of their application in metagenomics.

A few bioinformatics tools have been proposed to detect cross-sample contamination in genomic [13–15] or transcriptomic sequencing projects [16] without the need for controls or spike-ins. However, to the best of our knowledge, only one method has been developed specifically for metagenomics. Proposed by Lou et al., this approach tracks unexpectedly shared strains between samples processed together [5]. Few strains consistently found across multiple samples evenly distributed on a plate suggest external contamination, whereas many shared strains between physically close samples indicate cross-sample contamination. However, the method has several limitations. When two samples share multiple strains, identifying the contamination source and estimating contamination rates is challenging, especially if samples contaminate each other. Natural strain sharing, as observed in related individuals (vertical transmission) or cohabiting populations (horizontal transmission) [17, 18], further complicates the application of this method. Detecting shared strains requires *de novo* genome assembly and at least 50% genome coverage in the contaminated sample, reducing sensitivity for low-level contamination at low sequencing depths. Additionally, strain-level analyses are computationally intensive and time-consuming. Finally, despite promising results, the lack of an automated, user-friendly tool limits the broader adoption of this method.

Here, we introduce CroCoDeEL (CROss-sample COntamination DEtection and Estimation of its Level), a tool for accurate detection of cross-sample contamination. CroCoDeEL identifies both contamination sources and contaminated samples while estimating contamination rates. Designed as a decision-support tool, CroCoDeEL provides intuitive graphical reports, enabling users to interpret contamination events and perform manual curation if needed. A key advantage of CroCoDeEL is its reliance solely on species abundance profiles, which are fast and easy to generate. Unlike other approaches, it does not require negative controls or prior knowledge of samples location on the plate. CroCoDeEL implements a novel method that identifies specific patterns, which occur when comparing taxonomic profiles of a contaminated sample and its contamination source. CroCoDeEL is based on rule-based filtering, linear modeling, and a random forest model pre-trained on a human-curated semi-simulated dataset. To assess its performance, we tested CroCoDeEL on three independent metagenomic cohorts, demonstrating strong agreement with human experts classifications. Beyond validation, applying CroCoDeEL to published metagenomic datasets revealed extensive contamination, raising concerns about the robustness of conclusions in some high-impact studies. These findings underscore the urgent need for systematic cross-sample contamination screening, positioning CroCoDeEL as an essential tool for ensuring the reliability of sequencing data in metagenomics.

## Results

### Specific patterns in species abundance profiles are associated with cross-sample contamination

In theory, all the species from the contamination source should be introduced into the contaminated sample after a cross-sample contamination event. Some of these species could initially be absent from the contaminated sample before contamination. As cross-sample contamination can be thought of as a sample dilution, the relative abundances of these contamination-specific species are expected to be proportional between the contaminated and the contaminating sample; the proportionality coefficient being equal to the contamination rate. This insight forms the foundation of the CroCoDeEL approach.

To better illustrate this phenomenon, we compared species abundance profiles of a contaminated sample and its contamination source on a toy example (Fig. 1A). To do so, we visualized these profiles using a scatter plot with a logarithmic scale to enhance the visibility of subdominant species (Fig. 1B). Remarkably, this visualization reveals a subset of species characterized by a linear trend having a proportional abundance between the two samples (Fig. 1C). These are the contamination-specific species mentioned above, forming what we will subsequently call a contamination line.

**Fig. 1.**
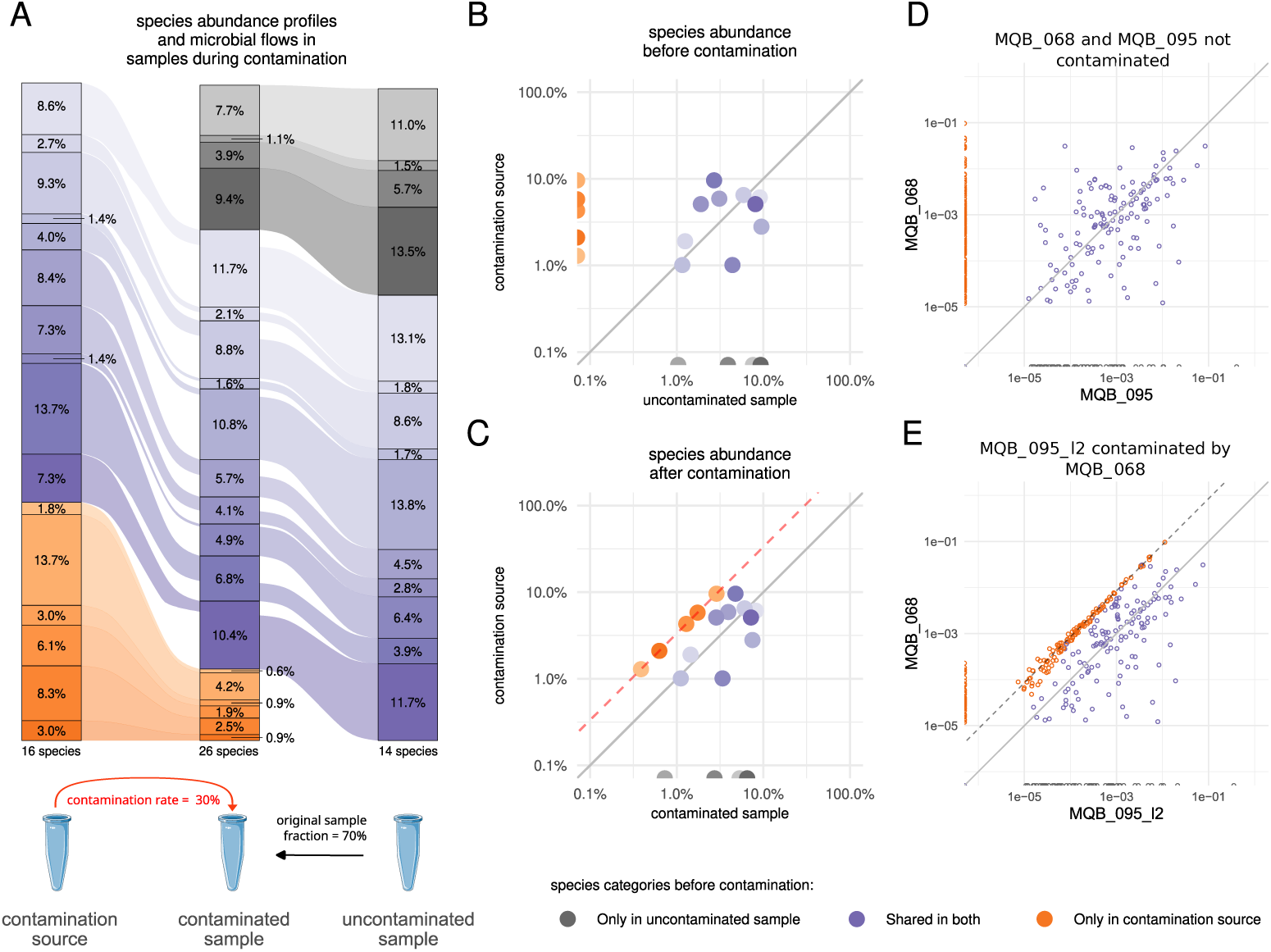
Illustration of cross-sample contamination patterns on a toy example and real metagenomic data. (A) Alluvial plot representing how contamination impacts species abundance profiles of a formerly non-contaminated sample. Before contamination, three species subsets can be distinguished: species shared by both the non-contaminated sample and the contamination source (blue), those specific to the non-contaminated sample (gray), and those specific to the contamination source (orange). (B) When comparing the species abundance profiles of the uncontaminated sample and its future contamination source on a log-transformed scatter plot, no distinct pattern emerges. Since log10(0) is undefined, species not detected are represented as half-points on either the x-axis or y-axis. (C) After contamination, abundances of contamination-specific species (orange dots) exhibit a noticeable linear pattern (red dashed line), with the equation y = x − log_10_(contamination rate) indicating proportionality. (D) Comparison of the species abundance profile of samples MQB 068 and MQB 095, which do not contaminate each other. (E) Comparison of the abundance profile of the same sample pair, but where MQB 095 l2 is intentionally contaminated by MQB 068 at a rate of approximately 10%.

Beyond this theoretical example, we performed an experiment to confirm that this phenomenon can be observed in real metagenomic data. Specifically, we collected two fecal samples, *MQB 068* and *MQB 095*, from two unrelated human individuals. After DNA extraction, we created a third sample, *MQB 095 l2*, by mixing *MQB 095* and *MQB 068* in approximately 90:10 proportions. In this context, *MQB 095 l2* represents *MQB 095* contaminated by *MQB 068* at a rate of 10%. The two original uncontaminated samples and the deliberately contaminated sample were subjected to shotgun metagenomic sequencing. As in the toy example, we first compared the species abundance profiles of *MQB 095 l2* and *MQB 068* using a log-scale scatterplot Fig. 1E. Two key observations confirmed *MQB 095 l2* as the contaminated sample with *MQB 068* as the source. First, above a certain abundance threshold, all species from *MQB 068* are detected in *MQB 095 l2*, as evidenced by the absence of points along the y-axis. This indicates that all species from *MQB 068* were introduced into *MQB 095 l2*, though species below the metagenomic detection threshold were not observed. Second, among the species shared between the two samples, some align along a straight line, representing species from *MQB 068* specifically introduced into *MQB 095 l2* through contamination. These species exhibit proportional abundance between the two samples, with higher abundance in *MQB 068*, as shown by the contamination line appearing above the identity line. In contrast, no contamination line is observed when comparing the species abundance profiles of the two uncontaminated samples (Fig. 1D). Furthermore, species richness in *MQB 095* is substantially lower than in its contaminated counterpart (194 vs. 319), indicating that 39% of the species detected in *MQB 095 l2* are artifacts of contamination.

Our goal was to develop a method capable of automatically detecting contamination lines, when present. To achieve this, we first created a semi-simulated dataset with species abundance profiles for 13,330 sample pairs, of which 44% were contamination events created by mixing data from real metagenomic samples originating from different projects. We then implemented an algorithm based on linear modeling to detect potential contamination lines within sample pairs. If a line was identified, ten expert-curated features were extracted (Fig. S1), including the number of species constituting the line and linear regression quality metrics. The algorithm was applied to all sample pairs in the semi-simulated dataset. Using these features, we trained a Random Forest classifier to distinguish contaminated from non-contaminated sample pairs. The model demonstrated excellent performance during validation, achieving precision and recall both exceeding 0.99. For each contamination event, the contamination rate was estimated by calculating the abundance ratio of species on the contamination line between the contaminated and the source sample. Provided that the reference used for taxonomic profiling is representative of species present in samples, this approach produced estimates closely matching theoretical values, with a median (Q1–Q3) absolute relative error of 8.3% (3.4%–16.9%) on the semi-simulated dataset. However, if abundant species in the contaminated sample or the contamination source are missing in the reference, the contamination rate can be respectively under- or overestimated (Fig. S2).

Finally, we developed CroCoDeEL, which combines both the feature extraction algorithm and the pre-trained model to classify all sample pairs from a real dataset as either contaminated or not, and estimate the respective contamination levels when relevant (Fig. 2).

**Fig. 2.**
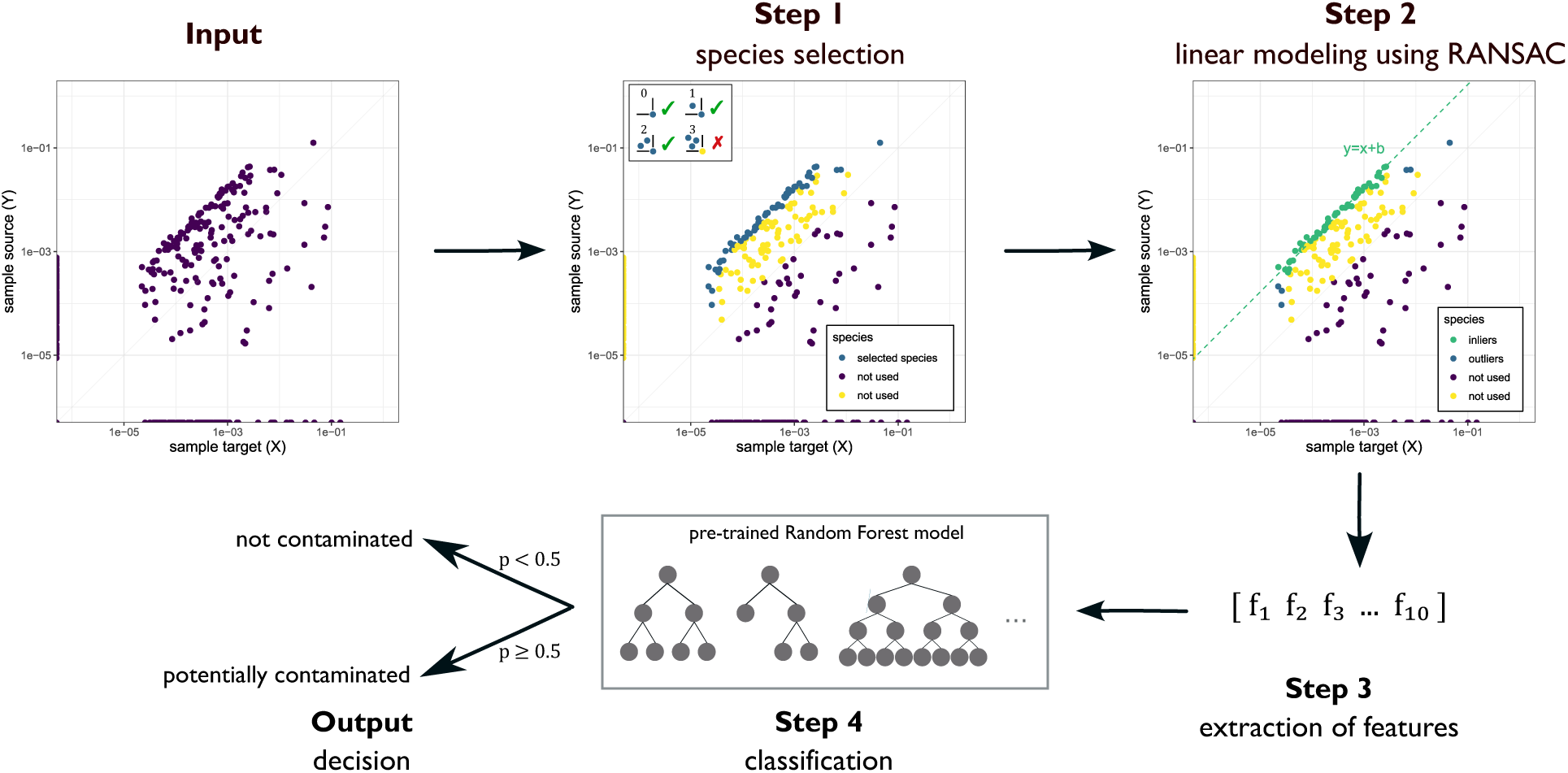
Flowchart describing the classification algorithm implemented in CroCoDeEL. Each step of the algorithm is detailed extensively in the Methods section. In this example, CroCoDeEL classifies a sample pair X and Y, where X is the suspected contaminated sample and Y is the suspected contamination source. **Input:** CroCoDeEL takes as input the log-transformed species abundance profiles of X and Y, which are visualized in a scatter plot. **Step 1:** Candidate species for a contamination line are selected. These species (blue points) have none or very few other species in their upper-left quadrant. **Step 2:** A potential contamination line is detected using the RANSAC regressor, which estimates the line offset (parameter b) and classifies candidate species as inliers (green points) or outliers (blue points). **Step 3:** Ten numeric features describing the contamination pattern are extracted. **Step 4:** The extracted features are input into a pre-trained Random Forest model for classification. **Output:** If the confidence score (probability) returned by the model exceeds 0.5, sample X is flagged as contaminated by sample Y, and the contamination rate is estimated based on the contamination line offset. Otherwise, sample X is considered as non-contaminated by sample Y.

### CroCoDeEL accurately identifies cross-sample contamination in real metagenomic data

To assess CroCoDeEL’s classification performance on real metagenomes, two human experts manually searched for cross-sample contamination events across three independent test datasets of human fecal samples. In each dataset, they meticulously inspected the species abundance profiles for all sample pairs to identify potential contamination lines, assigning a confidence level (low, medium, or high) to each suspected case. After consensus, these expert-labeled results served as the ground truth and were subsequently compared with CroCoDeEL classifications.

In each test dataset, the proportion of human-reported contamination events among all inspected sample pairs was low (*<* 0.5%). Therefore, unlike in the training dataset, real datasets tend to have a strong imbalance, with the majority of instances belonging to the non-contamination class. For this reason, we used performance indicators suited to this type of distribution [19, 20].

CroCoDeEL classification performance was consistent across the three datasets (Table 1, Supplementary Table 2). The Matthews correlation coefficient was around 0.7, indicating consistent classification accuracy for CroCoDeEL. The recall averaged 95%, meaning that the tool detected most of the contamination cases flagged by humans. Interestingly, among contamination cases reported by both CroCoDeEL and humans, we observed several cases where two contamination lines were visible on both sides of the identity line, representing scenarios in which samples contaminated each other (Fig. 3A).

**Fig. 3.**
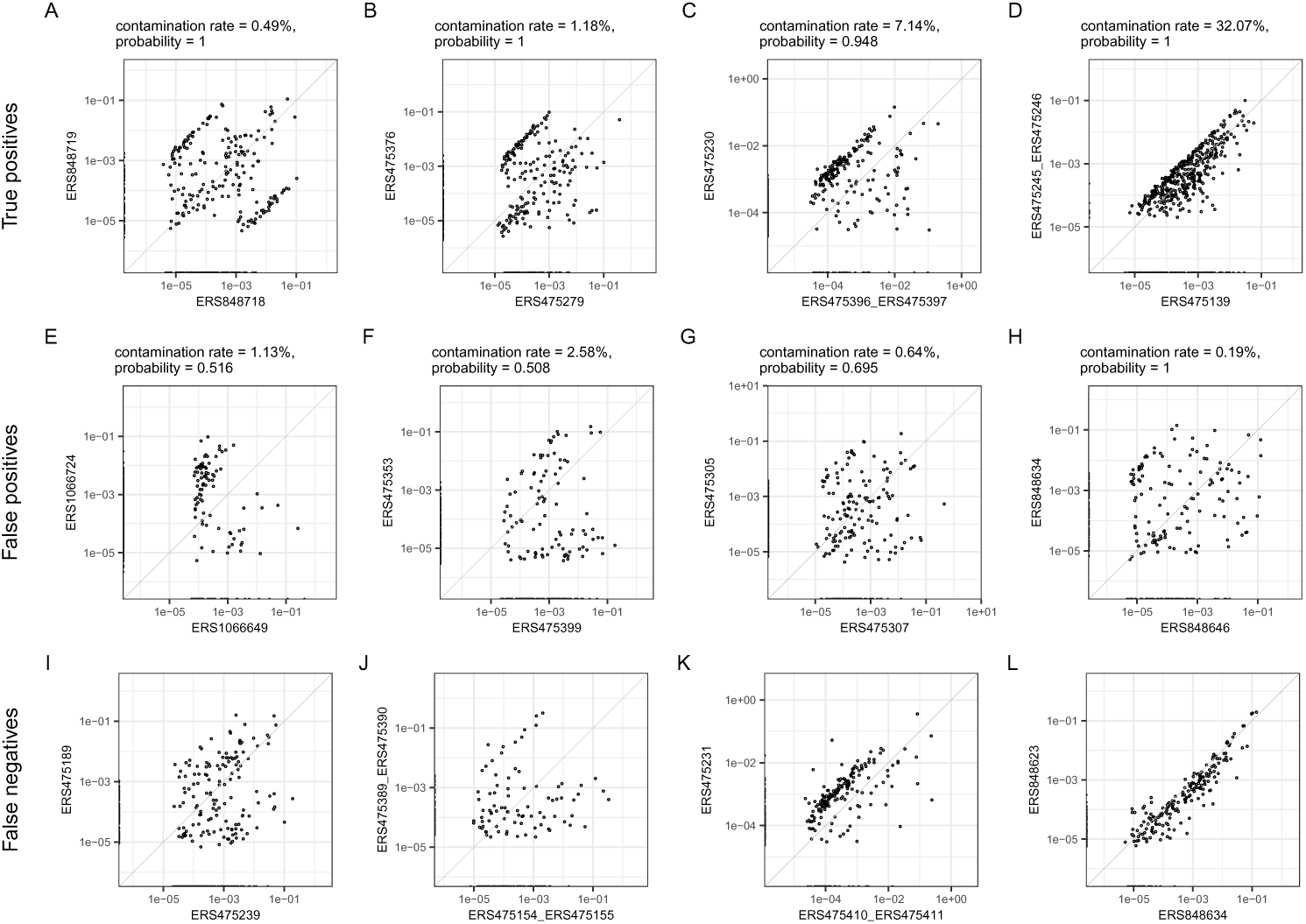
Scatter plots comparing on a logarithmic scale the species abundance profiles of sample pairs from the three test datasets. (A-D) Examples of contamination cases reported by humans and CroCoDeEL (i.e. true positives) (E-H) Examples of contamination cases reported by CroCoDeEL but not by humans (i.e. false positives) (I-L) Examples of contamination cases reported by humans but not by CroCoDeEL (i.e. false negatives)

**Table 1.**
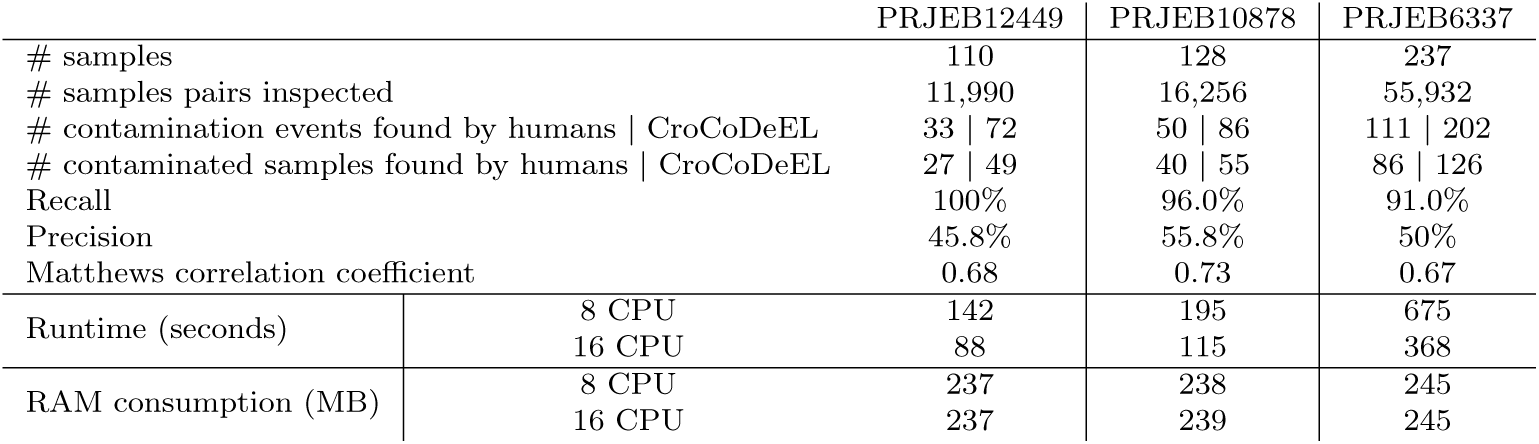
CroCoDeEL performance indicators on the three test datasets. Recall indicates the percentage of expert-identified contamination events found by CroCoDeEL, while precision indicates the percentage of cases reported by CroCoDeEL matching those of experts. Matthews correlation coefficient indicates the degree of association between CroCoDeEL predictions and expert classifications.

We identified two categories of false negatives. The first included contamination cases at very low rates, where human experts had limited confidence in their decision (Fig. 3I,J). The second, more concerning category, involved a few high-level contamination events with blurred contamination patterns, resulting in missed detections in cases we aimed to capture (Fig. 3K,L). For contamination case K, both implicated samples were contaminated by a third sample (*ERS475210 ERS475211*), albeit at different rates. As a result, the two samples shared a large number of species with proportional abundances. By transitivity, this led to the appearance of a blurred line when comparing their species abundance profiles, which, however, did not arise from direct contamination. We hypothesized that other cases where a third-party sample could not be identified may result from what we term “cascade contamination” where one sample contaminates another and is subsequently contaminated itself.

The specificity averaged 50%, indicating that half of the cases flagged by CroCoDeEL were not reported by humans (Fig. 3E,H). In each dataset, these potential false positives represented only about a hundred cases, or less than 0.5% of all sample pairs analyzed by CroCoDeEL. Most of these cases had low estimated contamination levels (average: 0.26%) and received a confidence score from the classification model significantly lower than that of true positives (mean probability: 0.73 vs. 0.92, p-value *<* 2.2 × 10^−16^, Mann–Whitney U test). Although a small proportion (*<*10%) of these events are likely to be false positives (Fig. 3E), the majority falls within a gray zone where contamination is plausible but at levels near the detection limit (Fig. 3F-H). This suggests that many of these ‘false positives’ may represent genuine, albeit minimal, contamination. To further test this hypothesis, we pooled all samples from the three datasets and classified with CroCoDeEL all sample pairs where cross-contamination was impossible, as the samples originated from distinct datasets. Among the 140,972 sample pairs considered, only five false positives were detected (Fig. S3), with reported contamination rates remaining low (mean = 0.28%). This result underscores the high specificity of CroCoDeEL in most scenarios where contamination is definitively absent.

In terms of computing resource utilization, CroCoDeEL searched for contaminations in a hundred samples and generated corresponding graphical reports within a few minutes on a standard computing node. Runtime was proportional to the number of sample pairs to be inspected, i.e. quadratic with respect to the number of samples. Additionally, CroCoDeEL demonstrated good scalability with the number of CPUs, achieving an efficiency of approximately 0.85. Its memory usage was low and remained relatively constant.

### CroCoDeEL performance is impacted by sequencing depth, contamination rates, and accuracy of species abundance profiles

Identifying a contamination line in metagenomic samples requires both the detection of contamination-specific species and their accurate quantification. This task becomes particularly challenging at low contamination rates or shallow sequencing depths, where contaminant species are typically subdominant and represented by a limited number of reads. To investigate how these factors influence CroCoDeEL’s performance, we simulated 25 sample pairs, with one sample in each pair contaminating the other. We varied both sequencing depth and contamination rates across different scenarios, generating species abundance tables for each condition using three different taxonomic profilers, Meteor2 [21], our in-house taxonomic profiler, along with two other popular tools, MetaPhlAn4 [22] and sylph [23]. These tables were then processed by CroCoDeEL, and the identified contaminations events were compared to the expected outcomes (Supplementary Table 3, Fig. 4).

**Fig. 4.**
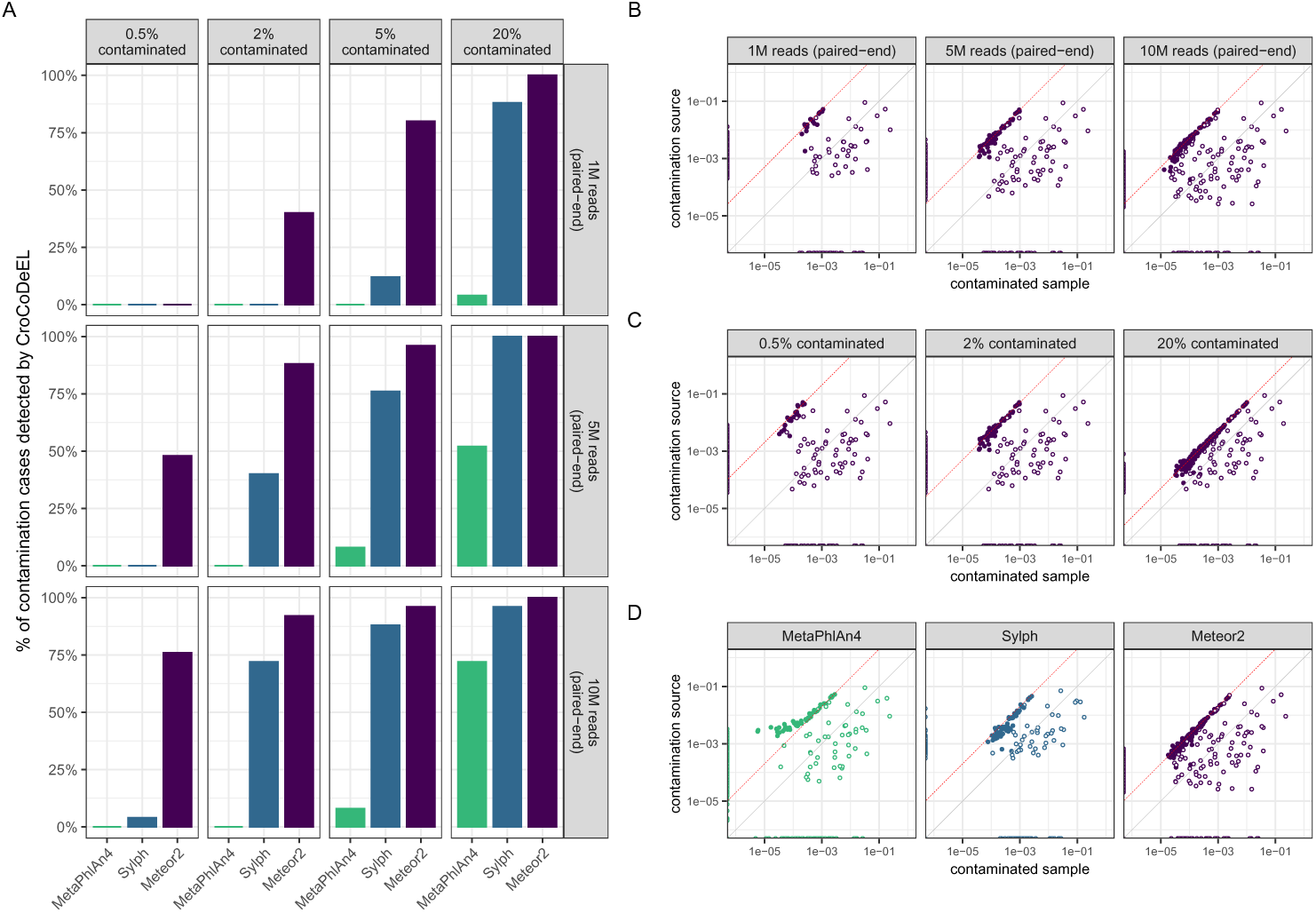
Impact of sequencing depth, contamination rate and taxonomic profilers on CroCoDeEL sensitivity. Performance of CroCoDeEL was evaluated using 25 semi-simulated sample pairs, assessing the influence of varying contamination rates (0.5%, 2%, 5%, 20%), sequencing depths (1M, 5M, 10M paired-end reads), and three taxonomic profilers (MetaPhlAn4, Sylph, Meteor2). (A) Proportion of contamination cases detected by CroCoDeEL (recall) for each profiler (colored bars) under different contamination rates (columns) and sequencing depths (rows). (B) Impact of sequencing depth: log-scale scatter plots comparing species abundance profiles of the same contamination case (2% contaminated, Meteor2 profiler) across varying sequencing depths. On all scatter plots, filled points indicate species introduced specifically by contamination. The theoretical contamination line is drawn in red. (C) Impact of contamination rate: log-scale scatter plots comparing species abundance profiles of the same contamination case (10M paired-end reads, Meteor2 profiler) under varying contamination rates. (D) Impact of taxonomic profilers: log-scale scatter plots comparing species abundance profiles of the same contamination case (10M paired-end reads, 5% contaminated) generated by the three taxonomic profilers.

These experiments confirmed that both sequencing depth and contamination rates are critical factors in detecting contamination events (p-values *<* 2 × 10^−16^, logistic regression). For instance, using Meteor2 profiles, CroCoDeEL successfully detected all cases of high contamination (rate = 20%), even with shallow sequencing (1M pairedend reads). In contrast, at a lower contamination rate (2%), CroCoDeEL detected 92% of contamination cases with 10M paired-end reads but only 40% with 1M paired-end reads (Fig. 4A). Higher sequencing depths increased the number of reads from contaminant species, enhancing their detection by taxonomic profilers and thus improving CroCoDeEL’s sensitivity (Fig. 4B). Similarly, higher contamination rates led to contaminant species comprising a larger proportion of total reads, further improving detection by taxonomic profilers and consequently boosting CroCoDeEL’s sensitivity (Fig. 4C).

In addition, our results demonstrated that CroCoDeEL’s sensitivity strongly depended on the profiler used (p-values *<* 2 × 10^−16^, logistic regression). Specifically, CroCoDeEL achieved much better sensitivity with species abundance profiles generated by Meteor2 than with those from Sylph or MetaPhlAn4, particularly at low sequencing depths and low contamination rates (Fig. 4A). For example, with 10M paired-end reads and a contamination rate of 0.5%, CroCoDeEL detected 76% of contamination cases with Meteor2, compared to only 4% with sylph and none with MetaPhlAn4. A comparison of the species abundance profiles generated by the three tools revealed that Meteor2 produced contamination lines comprising more species with less dispersion (Fig. 4D). Indeed, Meteor2 not only provided better sensitivity in detecting subdominant species introduced by contamination but also delivered more accurate quantification (Fig. S4). In contrast, MetaPhlAn4 systematically underestimated the abundance of subdominant species, which interfered with CroCoDeEL’s ability to detect contamination lines. Filtering out these species drastically improved CroCoDeEL’s sensitivity in some configurations (Fig. S5). However, Meteor2 was the only tool that produced abundance profiles suitable for effectively detecting contamination at low sequencing depths and/or low contamination rates.

### CroCoDeEL provides enhanced accuracy compared to strain-sharing-based methods

To our knowledge, aside from CroCoDeEL, only one other purely *in silico* method for detecting cross-sample contamination has been proposed. This approach, developed by Lou and colleagues [5], performs strain-level analysis and identifies contamination when unrelated samples share multiple strains. In their first case study, stool samples were collected from infants at multiple time points during their first year of life and from their mothers a few days after birth. A total of 402 samples were processed, with microbial DNA extracted in 96-well plates, followed by shotgun metagenomic sequencing. Samples were labeled using a family identifier and a suffix indicating the collection day for infants or ‘M’ for mothers. For example, *63D9* denotes a sample from the infant of family 63 collected on day 9, while *58M* refers to a sample from the mother of family 58. In this section, we focused on plate 3 (P3), where Lou and colleagues identified several cross-contaminated samples. We compared the results initially reported with those of CroCoDeEL after manual curation (Fig. 5, Supplementary Table 4). CroCoDeEL identified 16 human-validated contamination events involving 12 contaminated samples. In contrast, the strain-sharing-based method detected only two contaminated samples, both of which were also identified by CroCoDeEL. Notably, nine contamination cases identified by CroCoDeEL occurred between samples from adjacent wells, further highlighting the increased risk of contamination between physically close samples. Contamination sources were typically associated with older individuals, suggesting that well-to-well contamination, where low-biomass samples are contaminated by high-biomass samples or samples with greater microbial diversity, is more readily detected. This pattern reflects progressive diversification and increasing microbial load over time.

**Fig. 5.**
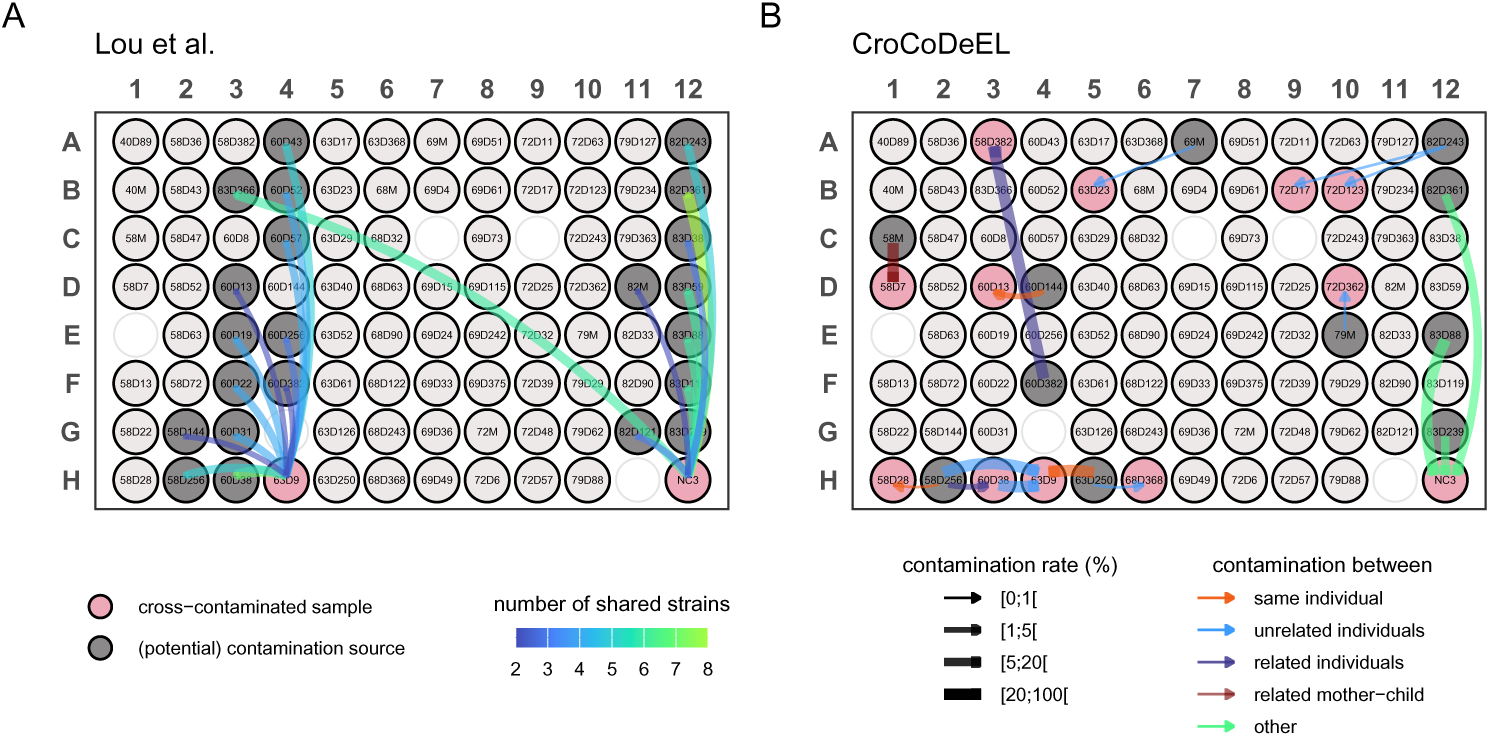
Comparison of contamination results on plate P3 using the strain-sharing method (Lou et al.) or CroCoDeEL. (A) Results from Lou et al.’s strain-sharing method. Pink wells represent contaminated samples, and gray wells indicate potential contamination sources. Arrow colors reflect the number of shared strains between the contaminated sample and the suspected sources. (B) Results from CroCoDeEL. Pink wells represent contaminated samples, and gray wells indicate contamination sources. Arrows show contamination events, with the direction pointing from the source to the contaminated sample. The thickness of the arrows corresponds to the estimated contamination rates, and their colors represent the relationships between samples.

Both tools identified two contaminated samples: the negative control *NC3* and *63D9*. According to the strain-sharing method, *63D9* was contaminated by samples from infants 60 and 58, while NC3 was contaminated by samples from infant 83 and members of family 82. However, since samples from the same individual or related individuals may naturally share strains, the exact contamination sources for these samples could not be determined using strain analysis alone. In contrast, CroCoDeEL not only identified the contaminated samples but also pinpointed the contamination sources and estimated contamination rates. For instance, CroCoDeEL revealed that sample *63D9* was specifically contaminated by *58D256* and *60D38* (Fig. 4A), which aligned with the observation that these samples shared the highest number of strains with *63D9* (7 and 5, respectively). A comparison of species abundance profiles for *63D9* and other potential contamination sources further confirmed these results (Fig. 4A). Similarly, CroCoDeEL determined that *NC3* was contaminated by samples *82D361*, *83D88*, and *83D249*, while excluding 82M, a sample from a mother, as a source (Fig. 4B).

Beyond these shared findings, CroCoDeEL identified additional contamination cases involving samples from related individuals or those collected from the same infant at different time points. Such cases were undetectable using the strain-sharing approach due to strain persistence in longitudinal data and vertical transmission between mothers and their children. However, CroCoDeEL successfully identified these events by accounting for the significant evolution of the infant microbiome over time and its distinct composition compared to adults (Fig. S6C). For instance, CroCoDeEL determined that sample *63D9* was strongly contaminated by 63D250, a sample from the same infant collected at a later time point. Similarly, sample *58D256* contaminated *60D38*, with infants 60 and 58 identified as twins. Regarding mother-to-infant contamination, CroCoDeEL identified a case in family 58 where sample 58D7 was contaminated by the mother’s sample, 58M. The species abundance profiles of these samples were highly similar (Spearman’s *ρ* = 0.88), and the microbial richness of *58D7* was unusually high for an infant of this age (223 species), confirming the high contamination rate estimated by CroCoDeEL.

An intriguing case reported by CroCoDeEL involved samples collected from twins at one year old (*60D382* contaminating *58D382*), representing one of the rare instances of contamination between samples in distant wells. Although the species abundance profiles of these two samples were highly similar (Spearman’s *ρ* = 0.87), which explains the high estimated contamination rate (*>*60%), no clear contamination line was observed (Fig. S6E), indicating that this case may represent a false positive. Notably, while the infants were raised together, such similarity in microbiota composition is unexpected, as shown by earlier samples where the similarity was much lower (Fig. S6E). These observations suggest a potential issue with sample collection or laboratory processing.

Finally, CroCoDeEL identified additional contamination events between unrelated samples, such as *82D243* contaminating *72D17* (Fig. S6D). While these events could theoretically have been detected by the strain-sharing approach, they were likely overlooked due to the low contamination rate(*<*2%), highlighting CroCoDeEL’s superior sensitivity in such scenarios.

### Undetected cross-sample contamination leads to erroneous results

Next, we examined another study by Lou et al. [24] that investigated microbial colonization in infants during their first year of life, analyzing the same dataset after excluding contaminated plates P3 and P4. For plates P1, P2, and P5, no contamination was found in the negative controls, nor any shared strains between unrelated samples, leading them to conclude that these plates were free from cross-sample contamination and suitable for analysis. However, using CroCoDeEL, we identified eight contamination events involving P2, including a critical case where the infant fecal sample 57D8 was contaminated by its mother’s fecal sample 57M at a rate of 23% (Supplementary Table 4). Although the authors reported that strains from mother 57 colonized her child’s gut at birth but did not persist, we suggest that this colonization never occurred. Instead, the detection of these strains in the newborn sample was likely an artifact of cross-sample contamination (Fig. 6A).

**Fig. 6.**
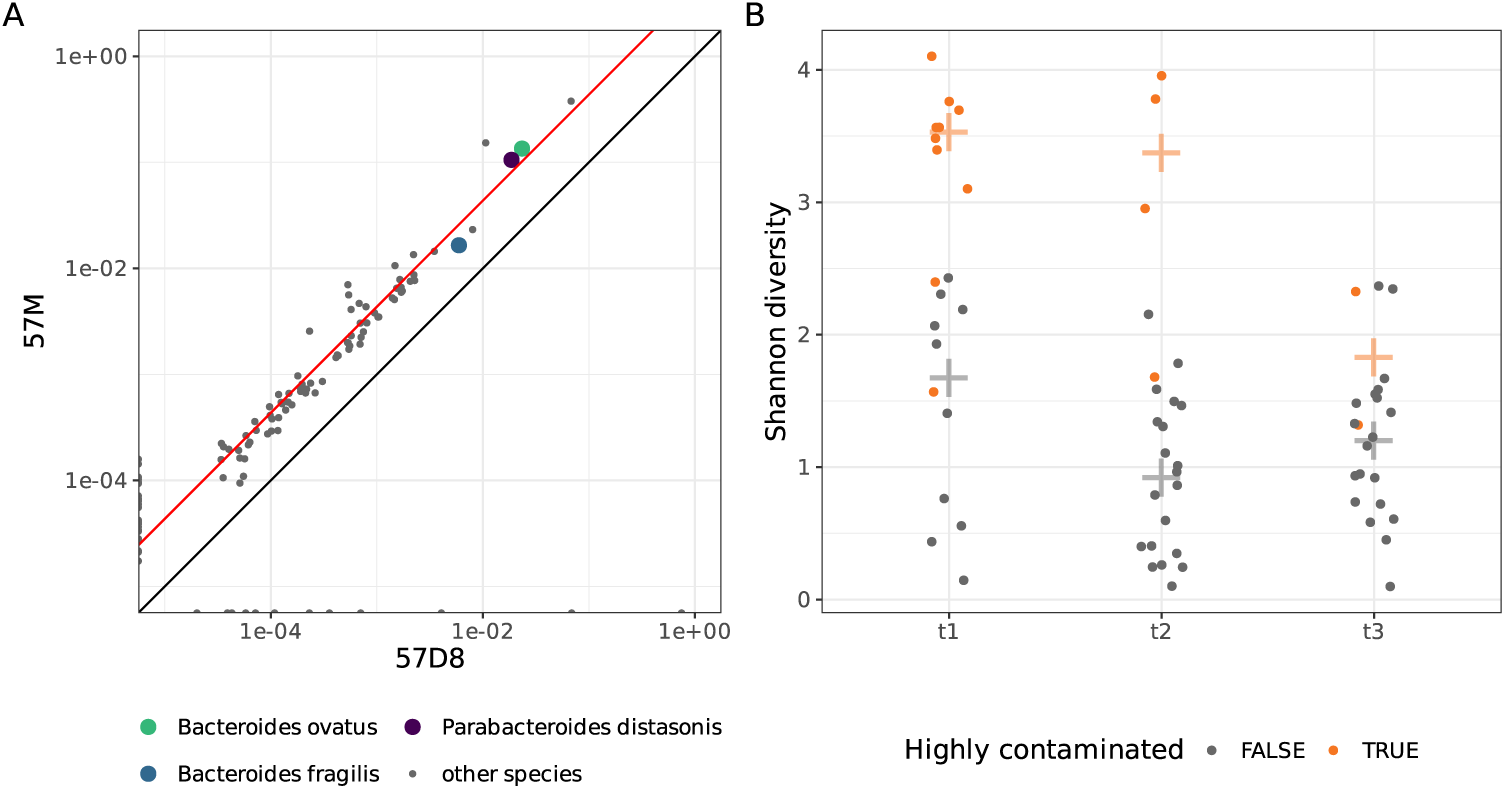
Cross-sample contamination affects studies combining high and low biomass samples. (A) Scatter plot comparing species abundance profiles between a newborn’s fecal sample (57D8, x-axis) and its mother’s (57M, y-axis). The infant sample is highly contaminated by the maternal sample, as shown by the red contamination line. Colored points highlight species identified by Lou et al. as non-persistent early colonizers but are likely contamination artifacts. (B) Strip charts comparing the Shannon diversity index in infant fecal samples from the Ferretti et al. study at three time points: t1 (1 day, n = 20), t2 (3 days, n = 24), and t3 (7 days, n = 22). Orange points indicate highly contaminated samples, while gray points represent other samples. Crosses indicate the median value for each sample type.

Similarly, a highly cited study by Ferretti et al. [25], which had a comparable purpose and design, encountered significant yet previously unreported issues with cross-sample contamination. CroCoDeEL identified 48 cross-contaminated samples out of 182 stool and tongue dorsum samples (Supplementary Table 5). In this case, 80% (45/56) of microbes reported as transiently present in some infants were artifacts of contamination, as they were found in newborn stool samples that were massively contaminated (10015, 10019, 10029, 10031, and 10055 at t1). As in the original study, we observed high species alpha-diversity in infant fecal samples at the first time point, followed by a significant decrease over the first week (p-value = 0.005 for t1 vs. t2; p-value = 0.003 for t1 vs. t3). However, due to cross-sample contamination, we argue that no reliable conclusions can be drawn. This apparent increase in diversity at t1 is primarily driven by highly contaminated samples. After their removal, no significant differences remain (p-value = 0.12 for t1 vs. t2; p-value = 0.56 for t1 vs. t3, Mann–Whitney U tests, Fig. 6B).

In a different context, we analyzed the TwinsUK cohort [26], which included 1004 fecal samples from adult donors. CroCoDeEL identified 202 contaminated samples, 176 of which were linked to the same eight sources of contamination (Supplementary Table 6). These eight sources exhibited highly similar species abundance profiles (Spearman’s *ρ >*0.96) and abnormally high species richness (mean: 782 ± 8), suggesting they were duplicated mixtures of multiple samples. Consequently, in many cases, the precise contamination source could not be determined. As most contaminated samples shared the same source, they were more similar to each other than uncontaminated samples were to each other (mean Bray-Curtis dissimilarity: 0.40 vs 0.53, p-value *<* 2 × 10^−16^, Mann–Whitney U test, Fig. 7A). Contaminated samples also exhibited significantly higher species richness (mean: 425 vs 303, p-value *<* 2 × 10^−16^, Mann–Whitney U test, Fig. 7B). Finally, 32% (440/1382) of species detected in at least 10 samples were more prevalent in the contaminated samples than in the uncontaminated ones (*FDR* ≤ 0.01, Fisher exact tests, Fig. 7C). These findings underscore the need to reassess the results of numerous studies based on this cohort, given the extent of the contamination problem.

**Fig. 7.**
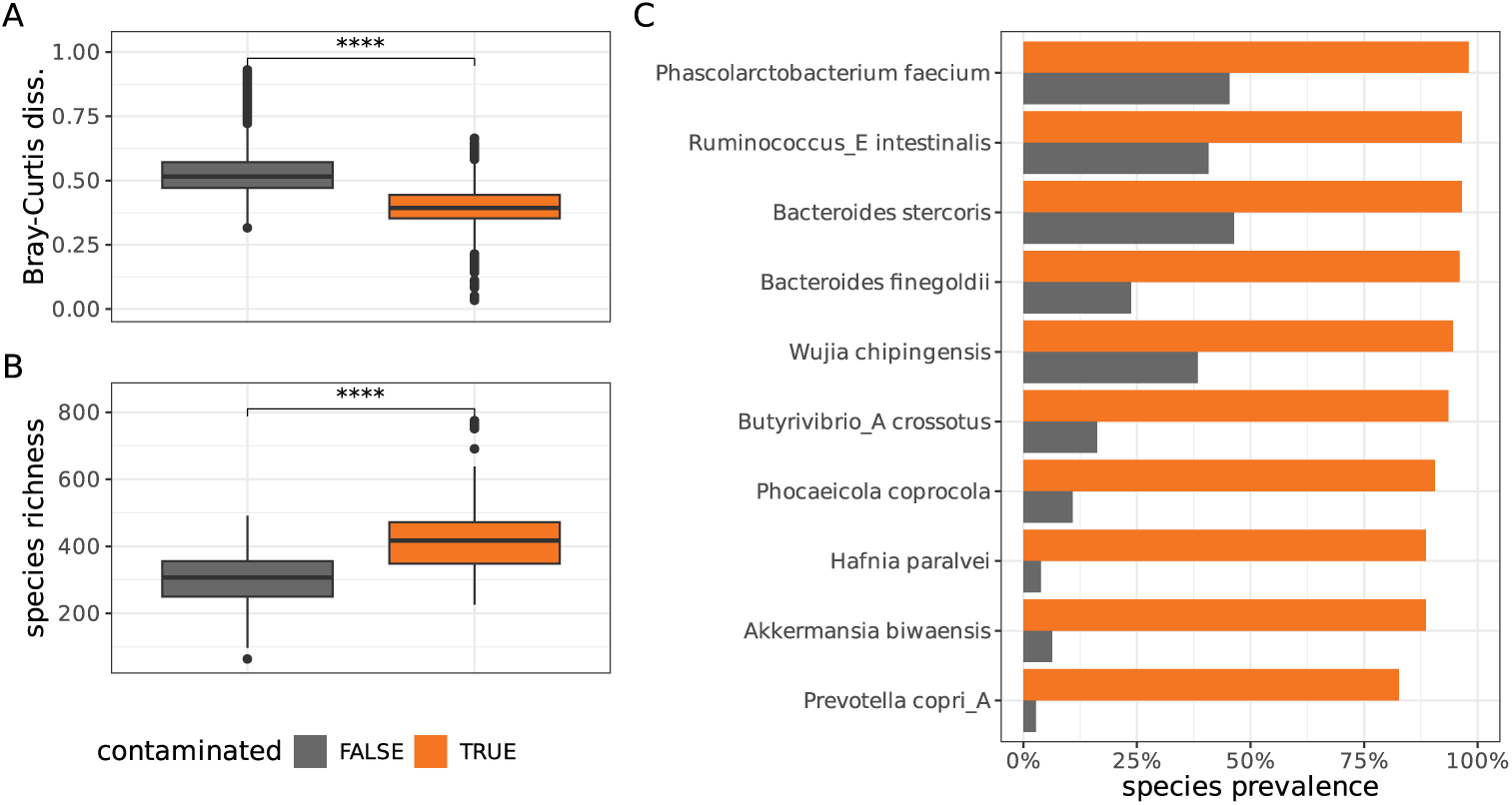
Cross-sample contamination affects the TwinsUK study. (A) Boxplots comparing the Bray-Curtis dissimilarity between all pairs of non-contaminated samples (n = 321,201; gray) and contaminated samples (n = 20,301; orange). (B) Boxplots comparing species richness between noncontaminated samples (gray, n = 802) and contaminated samples (orange, n = 202). (C) Bar plots showing the prevalence of species that are significantly more frequent in contaminated samples (orange) than in non-contaminated ones (gray). Statistical comparisons, when reported, were performed using Mann–Whitney U tests, with **** indicating a p-value < 2 *×* 10*^−^*^16^.

## Discussion

In this study, we demonstrate that cross-sample contamination distorts species abundance profiles and introduce a novel concept, the *contamination line*. This unique linear pattern emerges when comparing the species abundance profile of a contaminated sample with its contamination source. We verified that contamination lines are highly unlikely to occur by chance, typically appearing in fewer than one case per 10,000 sample pairs when analyzing human gut metagenomes. In the rare instances where they did occur, these lines involved only a few species, representing false positives at very low rates. By systematically searching for this contamination pattern across all sample pairs within a given cohort, we sought to identify contamination events as thoroughly as possible. Based on our experience, a trained human eye is highly effective at identifying such contamination lines. However, inspecting the abundance profiles of all sample pairs is highly time-consuming. For example, in a 96-well plate, identifying contamination would require reviewing 4,560 profiles. Given the rarity of contamination events, this process is prone to oversight. To streamline this task, we developed CroCoDeEL, an automated tool designed to detect contamination lines and minimize the likelihood of human error in large datasets.

A key advantage of CroCoDeEL is it reliance solely on species abundance profiles, eliminating the need for negative controls. This versatility enables seamless integration into existing workflows and ensures applicability across diverse metagenomic datasets, including those from public repositories. Additionally, CroCoDeEL does not require metadata about sample positions during laboratory processing and does not assume that cross-sample contamination is limited to adjacent samples on a plate. This flexibility allows for the detection of contamination between distant samples, including cases caused by leakage from a robotic pipetting arm or the dispersion of micro-droplets after removing a sealing film. Benchmarks based on real human fecal metagenomes confirmed that the tool detects cross-sample contaminations with high sensitivity provided sufficient sequencing depth. Notably, it was able to identify numerous likely cases at very low contamination rates that appeared negligible or were overlooked by human experts. Such sensitivity can be achieved only with taxonomic profilers that can detect and quantify subdominant species with high accuracy. However, this requirement is not adequately met by several popular taxonomic profilers. We propose that future efforts in developing and evaluating taxonomic profilers should place greater emphasis on accurately quantifying low-abundant species. In addition to contamination detection, CroCoDeEL estimates contamination rates, allowing researchers to evaluate the severity and impact of contamination on their datasets.

Despite its strengths, CroCoDeEL has limitations that will be addressed in future versions. First, as shown in real datasets, CroCoDeEL may occasionally miss complex contamination scenarios where contamination lines are less distinct. Additionally, the tool may falsely identify contamination in sample pairs with inherently similar species abundance profiles despite the absence of a contamination line. Such false positives are more likely in scenarios where samples are collected from animals raised in the same farm or cage environment or in longitudinal studies where samples are obtained from the same individual over short time intervals.

It is important to emphasize that CroCoDeEL is designed as a decision-support tool rather than a definitive classification system. Its purpose is not to replace human oversight but to significantly reduce the workload by identifying cases that warrant further review. Given the relatively small number of cases flagged by CroCoDeEL compared to the total number of sample pairs, human experts can efficiently confirm contamination by inspecting scatter plots in automatically generated reports and leveraging relevant metadata, if available.

Our findings suggest that cross-sample contamination is a prevalent issue in metagenomics, although its extent, both in terms of contamination rate and the proportion of affected samples, varies significantly across projects. We confirmed that it is particularly critical when mixing high and low-biomass samples, as low-volume contamination can lead to high DNA contamination. This highlights the need for caution when interpreting results from such experiments, as well as from followup analyses that reuse the data. For instance, several infant fecal metagenomes in public databases are contaminated with adult microbiota, potentially compromising Metagenome-Assembled Genome (MAG) collections of the early-life human gut microbiome [27, 28]. While the robustness of machine learning models and high statistical power in large cohort studies may help mitigate the impact of contamination-induced artifacts, the consequences are far more severe in clinical settings, where each sample is analyzed individually. Indeed, false discoveries of microbial species, including pathogens, can severely compromise diagnosis, prognosis, and risk assessments and other clinical applications relying on metagenomic sequencing. As this technology gains traction in medicine [29], the risks associated with cross-sample contamination require greater recognition.

CroCoDeEL’s ability to detect subtle cross-sample contamination as low as 0.1% raises the question of when contamination becomes problematic enough to warrant sample exclusion. The impact depends on study objectives and sample composition. In alpha diversity analyses, even a contamination rate as low as 1% can significantly inflate microbial richness, leading to the detection of dozens of spurious species. Conversely, since contaminants are typically present at low abundances, the Shannon diversity index, which accounts for taxa distribution, is only slightly affected.

Rather than relying solely on sample exclusion, addressing contamination may benefit from computational decontamination strategies. CroCoDeEL paves the way for such approaches, which aim to clean species abundance profiles by removing artifactual signals introduced by contamination. Although beyond the scope of this study, preliminary findings suggest that decontamination could serve as a valuable alternative to reprocessing samples. However, challenges remain, such as eliminating contamination-specific species without impacting naturally low-abundant taxa or addressing samples contaminated by multiple sources. Developing a comprehensive and robust decontamination tool will be a focus of future work, ensuring that metagenomic analyses are not only contamination-aware but also contamination-free.

As microbiome research scales up to include tens to hundreds of thousands of samples (e.g., the Microsetta Initiative, the Million Microbiome of Humans Project), increasing reliance on laboratory automation necessitates rigorous contamination monitoring. We therefore strongly advocate for integrating cross-sample contamination detection into routine sequencing quality control. By providing a robust framework for identifying such contamination, CroCoDeEL contributes to enhancing the reliability of microbiome research, clinical diagnostics, and other metagenomics-based applications.

## Materials and Methods

### Automatic detection of cross-sample contamination

This section details the algorithm implemented in CroCoDeEL, designed to automatically detect a contamination line, when present.

Let *X* = {*x*_1_*, x*_2_*, . . ., x_n_*} and *Y* = {*y*_1_*, y*_2_*, . . ., y_n_*} represent the abundances of *n* species in two samples, where *X* is the suspected contaminated sample and *Y* is the potential contamination source. Assuming contamination, we, define *r* ∈]0, 1] as the contamination rate. For species specifically introduced by contamination, we assume that 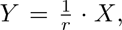 which, after log10 transformation, becomes log_10_(*Y*) = log_10_(*X*) −log_10_(*r*).

Subsequently, we compare log10 transformed values of *X* and *Y*. The algorithm is illustrated using a scatter plot, where *X* is plotted on the *x*-axis and *Y* (”Input” scatter plot, Fig. 2). Notably, this comparison is asymmetrical and must also be repeated by swapping the roles of *X* and *Y*. In this case, *Y* is the suspected contaminated sample, while *X* serves as the potential contamination source.

#### Step 1 Selection of candidate species for a contamination line

Contamination-specific species must be more abundant in the contamination source. Therefore, only species present in both *X* and *Y* but more abundant in *Y* are considered. Graphically, this corresponds to the set of points S*_upper_* above the identity line (blue and yellow points in the “Step 1” scatter plot, Fig. 2). When contamination occurs, the abundance ratio of the species shared between X and Y is supposed to be at least equal to the contamination rate. Graphically, this implies that no points should lie above the contamination line. Thereby, all species in S*_upper_* are screened and those with fewer than *l* = 2 other species in their upper left quadrant are selected. The resulting set of selected species is denoted as S*_candidates_* (blue points in the “Step 1” scatter plot, Fig. 2).

#### Step 2. Detection of the potential contamination line

If fewer than five candidate species are found, the sample pair is classified as uncontaminated, and the algorithm ends. Otherwise, the RANSAC (Random Sample Consensus) regressor [30] is applied to the points in S*_candidates_* to search for a contamination line. In our settings, this iterative algorithm fits a linear model *y* = *x* + *b*, where the estimated parameter *b* is the line offset. Points that fit the model are classified as inliers (S*_candidate inliers_*), while others are considered outliers (S*_candidate outliers_*). In the ‘Step 2’ scatter plot (Fig. 2), green points represent inliers, and blue points represent outliers. If fewer than five inlier species are detected, the sample pair is classified as uncontaminated, and the algorithm ends.

#### Step 3. Extraction of features describing the potential contamination line and its context

To verify that a detected contamination line represents a genuine contamination event, ten features identified as relevant by humans and selected using the Recursive Feature Elimination method (RFECV) are calculated. These features, listed below in order of decreasing predictive power, capture key characteristics of the contamination line and its context for downstream validation.(Fig. S5).

- (*f* 1) The number of species forming the contamination line (i.e number of inliers). A small number of inliers may suggest a spurious line.
- (*f* 2) The ratio between the number of species forming the contamination line and the number of species detected in both *X* and *Y*.
- (*f* 3) For each species forming the contamination line, the average Euclidean distance to the 5 nearest neighbors is calculated. The global average of these distances is then computed. This feature assesses the “compactness” of the contamination line.
- (*f* 4) For each species forming the contamination line, the average Euclidean distance to the 5 farthest neighbors is calculated, and the global average of these distances is computed. This feature assesses the “spread” of the contamination line.
- (*f* 5) The Spearman rank correlation coefficient between *X* and *Y*. A high Spearman’s rho between *X* and *Y* is expected at high contamination levels.
- (*f* 6) The mean orthogonal residual of inlier species to the contamination line, quantifying the line’s scatter.
- (*f* 7) The ratio between the number of species above the contamination line and the number of species detected in both *X* and *Y*.
- (*f* 8) The number of species above the contamination line. In the case of contamination, very few species should be above the line, except in complex scenarios such as cascade contaminations.
- (*f* 9) We begin by defining a pseudo-zero, which is one logarithmic unit below the smallest non-zero species abundance in the suspected contamination source (formally, *pseudo zero* = min (log_10_(*y_i_*)) − 1*, y_i_* ∈ *Y, y_i_ >* 0). Using this, we define an abundance cutoff *c*_1_ = *pseudo zero* + *b*, where *b* represents the estimated offset of the contamination line described earlier. We also define *m* as the average abundance in the source of the 10 most abundant species detected in the source but absent in the suspected contaminated sample (i.e., points on the y-axis). Finally, the feature |*m* − *c*_1_| quantifies whether highly abundant species in the potential source are unexpectedly missing from the contaminated sample.
- (*f* 10) We define another abundance cutoff *c*_2_ defined as average abundance in the source of the 10% least abundant inlier species (i.e, those forming the contamination line). The final feature, |*m* − *c*_2_|, serves the same purpose as *f* 9.

#### Step 4. Random Forest classification model to confirm a contamination event

The ten extracted features are fed into a pre-trained Random Forest model (1,000 trees) for classification. This classifier was trained using 13,350 sample pairs from the semi-simulated dataset, with labels indicating whether each pair was contaminated.

If the predicted probability of contamination is ≥ 0.5, the sample pair is classified as contaminated with a contamination rate *r* equal to 10^−*b*^. The algorithm then terminates, and this probability presents the confidence level of the classification. Conversely, if the probability is *<* 0.5, the sample pair is classified as not contaminated.

### Metagenomic data preprocessing and taxonomic profiling

For Illumina sequencing data, quality control was performed using fastp [31], including the following steps: 1) removal of sequencing adapters, 2) trimming or discarding low-quality reads, and 3) discarding reads shorter than 60 bp. For Ion Torrent sequencing data, quality control was performed using AlienTrimmer [32]. Reads mapped to the human genome (T2T CHM13v2.0, GCA 009914755.4) with Bowtie2 [33] were subsequently removed.

Unless otherwise specified, taxonomic profiling was conducted using the Meteor software suite [21]. For each sample, high-quality microbial reads were mapped with Bowtie2 [33] to the updated Integrated Gene Catalogue of the human gut microbiota (IGC2), comprising 10.4 million genes [34]. Alignments with nucleotide identities below 95% were discarded, and gene counts were computed using a two-step procedure previously described, which handles multi-mapped reads [35]. Raw gene counts were normalized according to gene length.

The IGC2 gene catalog has been clustered into 1,990 MetaGenomic Species (MGS) (clusters of more than 100 co-abundant genes belonging to the same microbial species) using MSPminer [36, 37]. The abundance of an MGS was calculated as the mean abundance of its 100 signature genes (i.e., the genes most correlated within the cluster). If fewer than 10% of the signature genes were detected in a sample, the species abundance was set to 0.

In addition to Meteor, taxonomic profiling was performed using Sylph (version: v0.6.0) [23], based on the Genome Taxonomy Database r214 [38], or MetaPhlAn4 (version: 4.1.0) [22], using the vJun23 database.

### Creation of real cross-contaminated samples

Nine fecal samples were self-collected by volunteers and immediately preserved in a stabilizing solution (Zymo DNA/RNA Shield). DNA extraction was performed by the SAMBO platform (MetaGenoPolis, INRAE, Jouy-en-Josas, France) using a semiautomated protocol based on SOP 07 V2 of the IHMS project [39, 40]. Contamination was simulated by mixing purified DNAs. Shotgun metagenomic sequencing was conducted at the MetaQuant platform (MetaGenoPolis, INRAE, Jouy-en-Josas, France) using Ion Torrent (Thermo Fisher Scientific, Waltham, USA) or DNBSEQ-G400 (MGI Tech, China) instruments as previously described [41]. A description of the newly sequenced samples is available in Supplementary Table 1.

### Creation of semi-simulated cross-contaminated samples

To mimic a cross-contaminated sample, contents from two real metagenomic samples were mixed in proportions corresponding to the expected contamination rate. Two strategies described below were used to achieve this.

Let *r* ∈]0, 1] and *n* ∈ N^∗^. Consider a simulated sample *C* consisting of *n* reads, where sample *A* is contaminated by sample *B* at a rate *r*.

1. Read-based subsampling: Reads were subsampled from the original samples using the ‘sample’ command implemented in seqtk (version: 1.4 r122) [42]. Specifically, *n* · (1 − *r*) reads were drawn from sample *A*, and *n* · *r* reads from sample *B*. The subsampled reads were merged into a new FASTQ file, representing the contaminated sample *C*. This file was then processed to generate taxonomic profiles, as described earlier.
2. Gene count-based rarefaction: Gene count vectors for samples *A* and *B* were first generated using Meteor [21]. For each sample, a pseudogene was added to represent unmapped reads. Rarefaction was performed by randomly selecting *n* · (1 − *r*) counts from *A* and *n* · *r* counts from *B*. The rarefied gene counts were summed to create a composite gene count vector for the contaminated sample *C*, which was subsequently analyzed with Meteor to produce species abundance profiles.

Both strategies successfully generated simulated samples with species abundance profiles resembling those of real cross-contaminated datasets (Fig. S1). The gene count-based approach is computationally faster but limited to generating species abundance profiles via Meteor. Conversely, the read-based subsampling method is more versatile, allowing comparisons across various taxonomic profilers, albeit at the cost of generating larger intermediate files.

An alternative approach using linear combinations of species abundance profiles was tested but deemed unsuitable due to two major shortcomings: it produced unrealistic sensitivity for detecting subdominant species introduced through contamination and produced an overly idealized contamination profile, with data points aligning perfectly along the contamination line (Fig. S7E-F).

### Generation of the semi-simulated training dataset

To train a classifier that will identify contamination events, we created a semi-simulated dataset with species abundance profiles for 15,000 sample pairs. To do so, eleven independent cohorts, including two with restricted access, were selected, comprising a total of 15,203 samples (Table 2). These cohorts were chosen to represent two microbial ecosystems (oral and human gut microbiota) and a broad range of phenotypes and geographical origins. For these samples, raw gene count tables (i.e., the number of mapped reads for each gene) were generated using the Meteor software suite, as described above.

**Table 2.** Description of the eleven cohorts used to generate the semi-simulated training dataset.

| BioProject acc. | Project name | Body site | Country | # samples | Reference |
| --- | --- | --- | --- | --- | --- |
| PRJEB37249 | MetaCardis BMIS | Human gut | EU | 888 | [43] |
| PRJEB32731 | Twins UK | Human gut | UK | 1004 | [44] |
| PRJEB11532 | Personalized nutrition glycemic response | Human gut | ISR | 969 | [45] |
| PRJEB38742 | MetaCardis Imidazole propionate | Human gut | EU | 1018 | [46] |
| PRJNA398089 | HMP2 IBD | Human gut | USA | 1338 | [47] |
| PRJNA354235 | Health Professionals Follow-up Study | Human gut | USA | 929 | [48] |
| PRJDB4176 | CRC | Human gut | JPN | 645 | [49] |
| PRJEB51353 | SCAPIS | Human gut | SWE | 3992 | [50] |
| PRJEB45799 | Oral transmission | Human oral | USA | 1929 | [51] |
| EGAD00001006554 | Milieu intérieur | Human gut | FR | 1356 | [52] |
| EGAD00001001991 | Lifelines deep | Human gut | NL | 1135 | [53] |

Initially, sample pairs were carefully selected from independent cohorts to ensure that they could not contaminate each other prior to simulation. Contamination was then simulated in half of the dataset (7,500 sample pairs) by mixing rarefied gene counts of the two samples, as explained above. Contamination rates were uniformly sampled from 0% to 100% for 5,500 sample pairs, and from 0% to 5% for the remaining 2,000 pairs. This dual sampling approach was designed to enhance the classifier performance, as empirical observations suggested that most contaminations occur at low levels (below 5%). Sequencing depth was varied across all sample pairs, contaminated and non-contaminated, by performing additional gene count rarefaction. Sequencing depth was uniformly sampled between 1 and 20 million reads. Finally, the gene counts of these simulated samples were processed through Meteor to generate species abundance profiles, as described in the previous subsections.

Species abundance profiles for each sample pair corresponding to contamination cases were visually inspected using scatter plots by human experts (L.G. and G.G.). Sample pairs were discarded if no contamination line was visible due to low sequencing depth or insufficient contamination rates. The final curated dataset comprised 7,480 non-contaminated sample pairs and 5,850 contaminated pairs.

### Generation of the test datasets

#### Real metagenomic datasets labeled by experts

Metagenomic sequencing data from three public cohorts were downloaded Table 3 and species abundance tables were generated with Meteor2 [21]. For each dataset, scatter plots of species abundance profiles were created, representing pairwise comparisons across all samples. These plots were systematically examined to identify potential contamination lines. When a contamination pattern was observed, both the contaminated sample and the contamination source were annotated. In total, 84,178 plots were reviewed independently by two evaluators (G.G. and L.G.).

**Table 3.** Description of the three cohorts manually curated by human experts to generate test datasets.

| Bioproject acc. | Body site | Country | # samples | # inspected scatter plots | Cohen's $\kappa$ | Reference |
| --- | --- | --- | --- | --- | --- | --- |
| PRJEB12449 | gut | USA | 110 | 11,990 | 0.78 | [54] |
| PRJEB10878 | gut | DNK and CHN | 128 | 16,256 | 0.76 | [44] |
| PRJEB6337 | gut | CHN | 237 | 55,932 | 0.52 | [35] |

Inter-rater reliability was assessed using Cohen’s *κ* coefficient [55], commonly used in psychological and behavioral studies. The *κ* values calculated for the three datasets (Table 3) highlight the challenges in achieving consensus for detecting contamination at low rates or in complex scenarios. Following the guidelines of Landis and Koch [56], *κ* values between 0.4 and 0.6 indicate moderate agreement, while values between 0.6 and 0.8 represent substantial agreement. These results underscore the inherent complexity of manually detecting cross-sample contamination, even for experienced evaluators. Discrepancies between the annotations by G.G. and L.G., which primarily arose from oversight, ambiguous cases, or typographical errors, were resolved through arbitration sessions involving both evaluators and a third reviewer (F.P.O.). During these sessions, contamination cases were assigned a confidence level (Low, Medium, or High) based on consensus (Supplementary Table 2). Notably, the arbitration process adopted a stringent criteria for confirming contamination, ensuring robustness in contamination labeling but potentially increasing the rate of false negatives.

#### Semi-simulated dataset to evaluate the impact of sequencing depth, contamination rates and taxonomic profilers

Metagenomic sequencing data from the bioprojects PRJNA763023 and PRJDB4176 were downloaded. 25 sample pairs were randomly selected, ensuring that each pair consisted of samples from different cohorts to prevent cross-sample contamination prior to contamination. Within each pair, one sample was designated as the contamination source and the other as the contaminated sample. Contamination was simulated by mixing reads between paired samples, as described earlier. Sequencing depth was varied by subsampling reads. The resulting dummy FASTQ files were then processed using Sylph, MetaPhlAn4, and Meteor2 to generate species abundance tables. CroCoDeEL classification results are available in Supplementary Table 3.

#### First case study by Lou et al

Metagenomic sequencing data from the bioproject PRJNA698986 were downloaded and a species abundance table was generated with Meteor2. CroCoDeEL classification results were curated by L.G, G.G. and E.L.C (Supplementary Table 4). Sample metadata and results of the strain analysis were obtained from the Yue Clare Lou personal GitHub repository.

### Implementation and benchmarks

CroCoDeEL was implemented in Python3. Species abundance tables and other data frames were managed with the Pandas module [57]. Implementation of the Random Forest and and the RANSAC algorithms available in the scikit-learn module [58] were used. Scatter plots comparing species abundance profiles of samples were generated using the Matplotlib module [59]. Finally, the multiprocessing module was used to process all the pairs of samples to be classified in parallel. All results reported in this study were obtained with CroCoDeEL version 1.0.3. Benchmarks were performed on a computing node with two Intel®Xeon®CPU E5-2680 clocked at 2.70GHz, each containing 8 physical cores, along with 256GB of RAM. RAM consumption and execution times were measured with GNU time.

**Supplementary information.**

## Declarations

### Data availability

Metagenomic sequencing data of the cross-contaminated samples generated for this study were deposited in the European Nucleotide Archive (ENA) under Bio-Project accession number PRJEB83730. Other public metagenomic and amplicon sequencing data are available on the INSDC databases through the bioprojects PRJDB4176, PRJEB10878, PRJEB11532, PRJEB12449, PRJEB32731, PRJNA352475, PRJEB37249, PRJEB38742, PRJEB45799, PRJEB51353, PRJEB6337, PRJNA398089, PRJNA698986, PRJNA354235, PRJNA763023, and PRJNA394849. Metagenomic sequencing data with restricted access are available on the European Genome-phenome Archive (EGA) after filling in a request form through the datasets: EGAD00001001991 and EGAD00001006554. Other data including species abundance profiles from training and testing datasets are available on the Recherche Data Gouv repository (dataset N6JSHQ).

### Software availability

CroCoDeEL source code is publicly available on GitHub under the GNU GPL v3 licence (https://github.com/metagenopolis/CroCoDeEL). Software documentation, including installation procedure, is also available on this repository.

## Acknowledgements

This work was funded by the MetaGenoPolis grant ANR-11-DPBS-0001. We thank Florence Thirion and Victoria Meslier for the in-depth testing of CroCoDeEL and their valuable feedback. We also thank Patrick Veiga for reviewing the draft version of this paper and providing helpful suggestions.

## Author contributions

E.L.C. designed the original method for graphically identifying contamination events. L.G. and G.G. developed the algorithm for automatic contamination detection. A.F. and B.Q. created the contaminated samples and performed shotgun sequencing. L.G., E.L.C., F.P.O., and G.G. designed and conducted the analyses. F.P.O. and L.G. wrote the software. G.G., E.P., E.B., F.P.O., and E.L.C. supervised the project. E.L.C., F.P.O., and G.G. wrote the manuscript. All authors read and approved the final manuscript.

## Ethics declaration

The study protocol involving collection of stool samples was approved by the French Committee for the protection of persons (CPP Ile-de-France 10) registered under reference 2021-A02873-38. Written consent was obtained from all participants. The authors declare no competing interests.

## Supplementary Tables

- Supplementary Table 1: Details of newly sequenced samples, including uncontaminated and intentionally contaminated cases.
- Supplementary Table 2: Classification results of human experts and CroCoDeEL on the three cohorts consisting of real human fecal metagenomes.
- Supplementary Table 3: CroCoDeEL results on the 25 semi-simulated sample pairs by varying the contamination rate, sequencing depth, and the taxonomic profiling tool.
- Supplementary Table 4: Contamination cases detected by CroCoDeEL on plates P2 and P3 of the Lou et al. dataset (PRJNA698986).
- Supplementary Table 5: Contamination cases detected in the Ferretti et al. dataset (PRJNA352475).
- Supplementary Table 6: Contamination cases detected in the TwinsUK dataset (PRJEB32731).

## Supplementary Figures

**Fig. S1.**
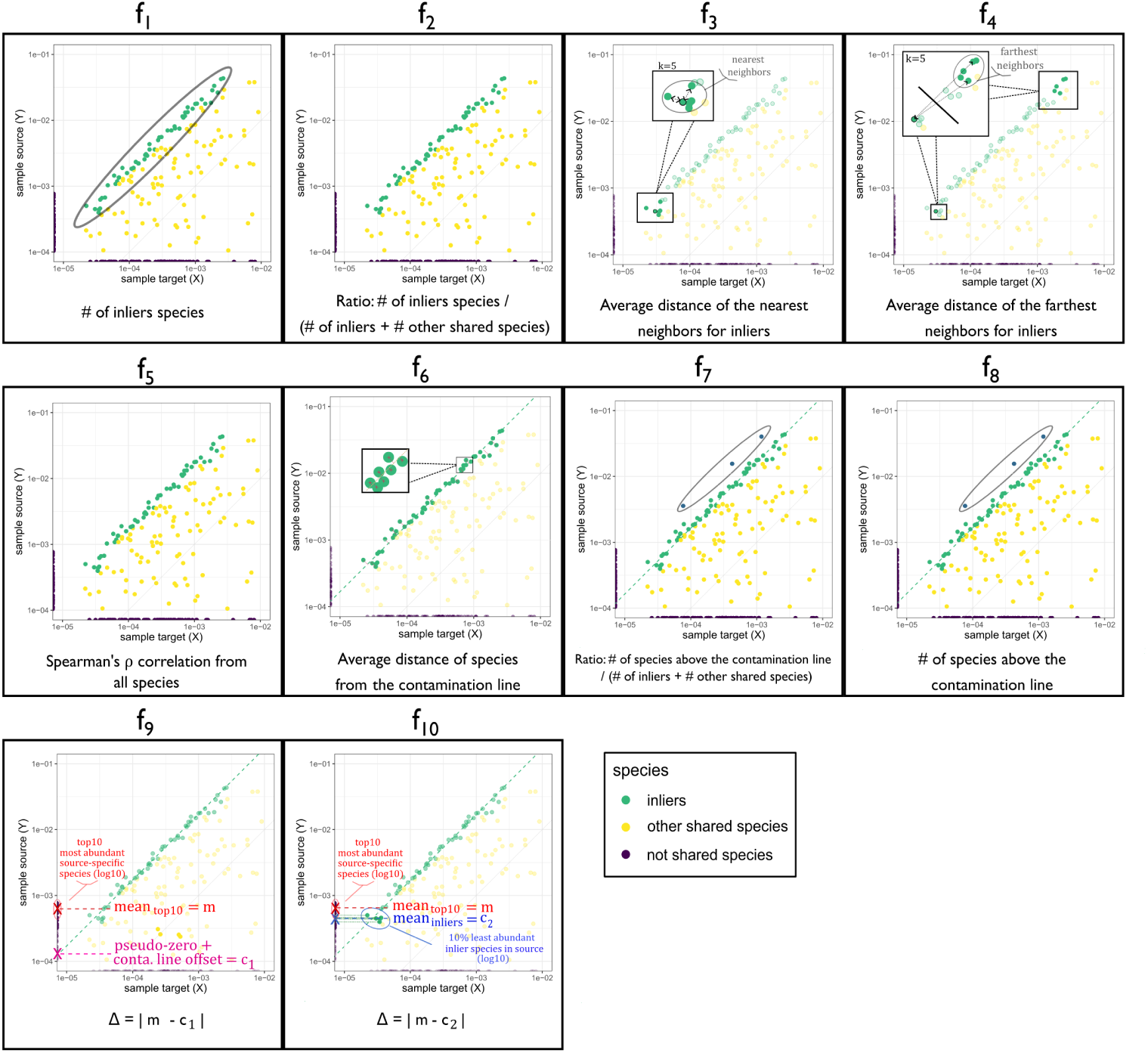
Illustration of ten features used by the Random Forest model to classify a sample pair as contaminated or not. Scatter plots show the species abundance profiles on a logarithmic scale, with the contamination source represented on the y-axis and the potentially contaminated sample on the x-axis. A detailed description of these features used in the step 4 of the classification algorithm is provided in the Methods section.

**Fig. S2.**
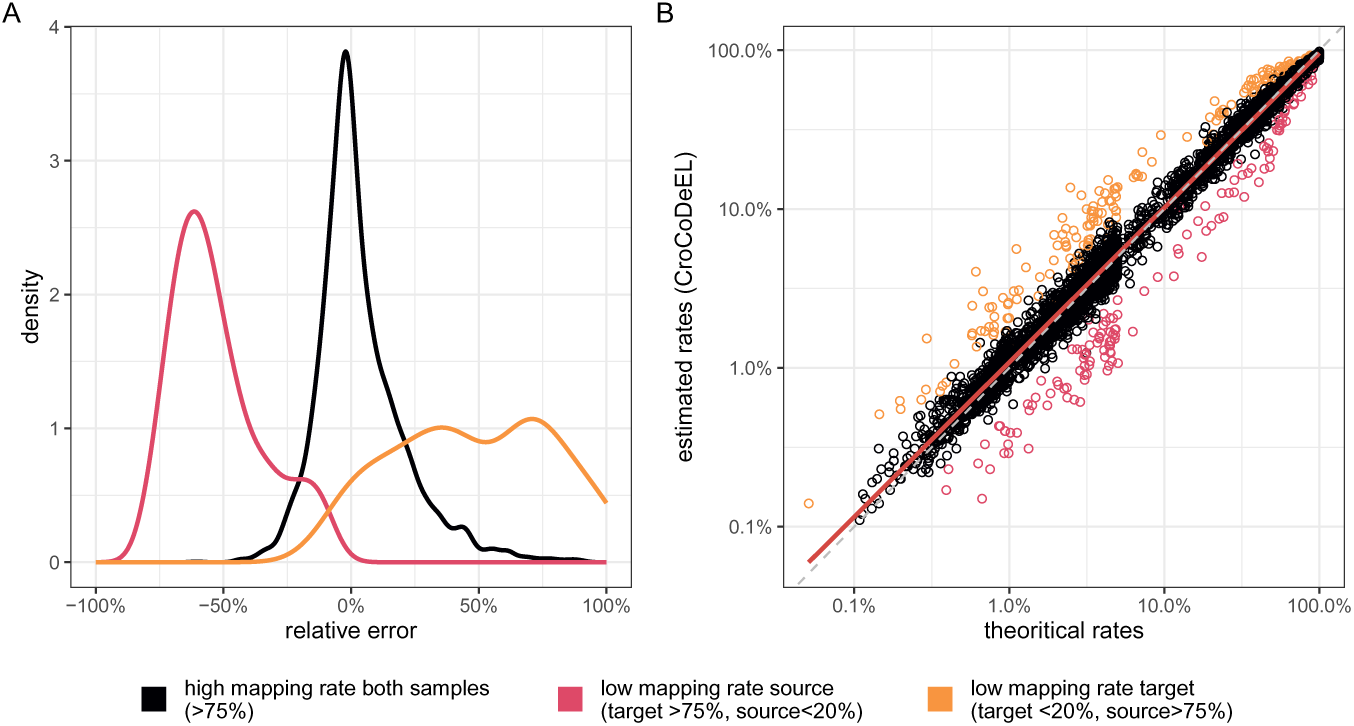
Evaluation of contamination rate predictions by CroCoDeEL. (A) Density plots depict the distribution of relative errors between estimated and theoretical contamination rates. (B) Scatter plot (log scale) of estimated versus theoretical contamination rates. Contamination rates estimated by CroCoDeEL were evaluated using 5,850 contaminated sample pairs from the semi-simulated training dataset. The analysis was conducted under three mapping conditions: (1) high mapping rates for both source and target samples (black), (2) low mapping rate for the source sample (target >75%, source <20, red) and (3) low mapping rate for the target sample (target <20%, source >75%, orange). CroCoDeEL reliably estimates contamination rates when both source and target samples exhibit high mapping rates. However, low mapping rates for the source sample lead to underestimation, while low mapping rates for the target sample result in overestimation.

**Fig. S3.**
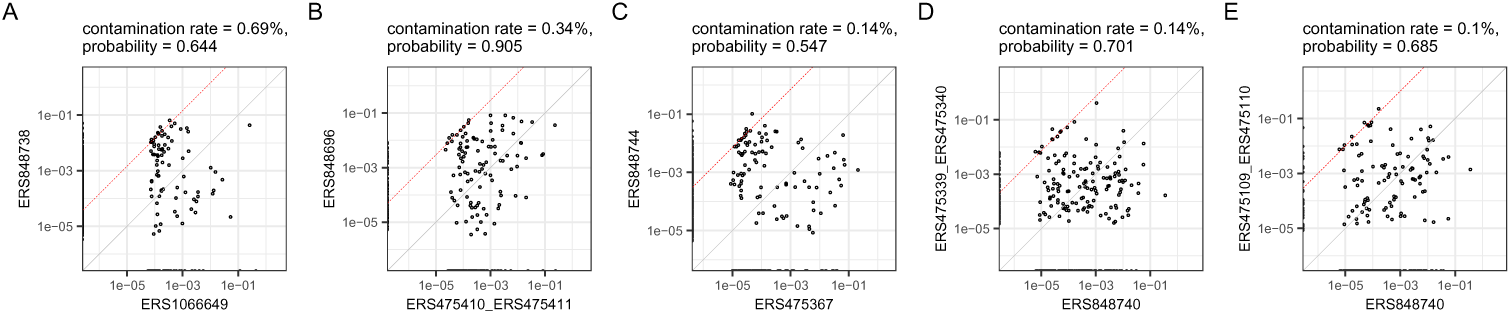
False positives reported by CroCoDeEL in non-contaminated sample pairs. CroCoDeEL analyzed 140,972 sample pairs where contamination was not possible and identified only five false positives. Log-scale scatter plots illustrate species abundance profiles for each false positive, along with the estimated contamination rate and probability returned the Random Forest model.

**Fig. S4.**
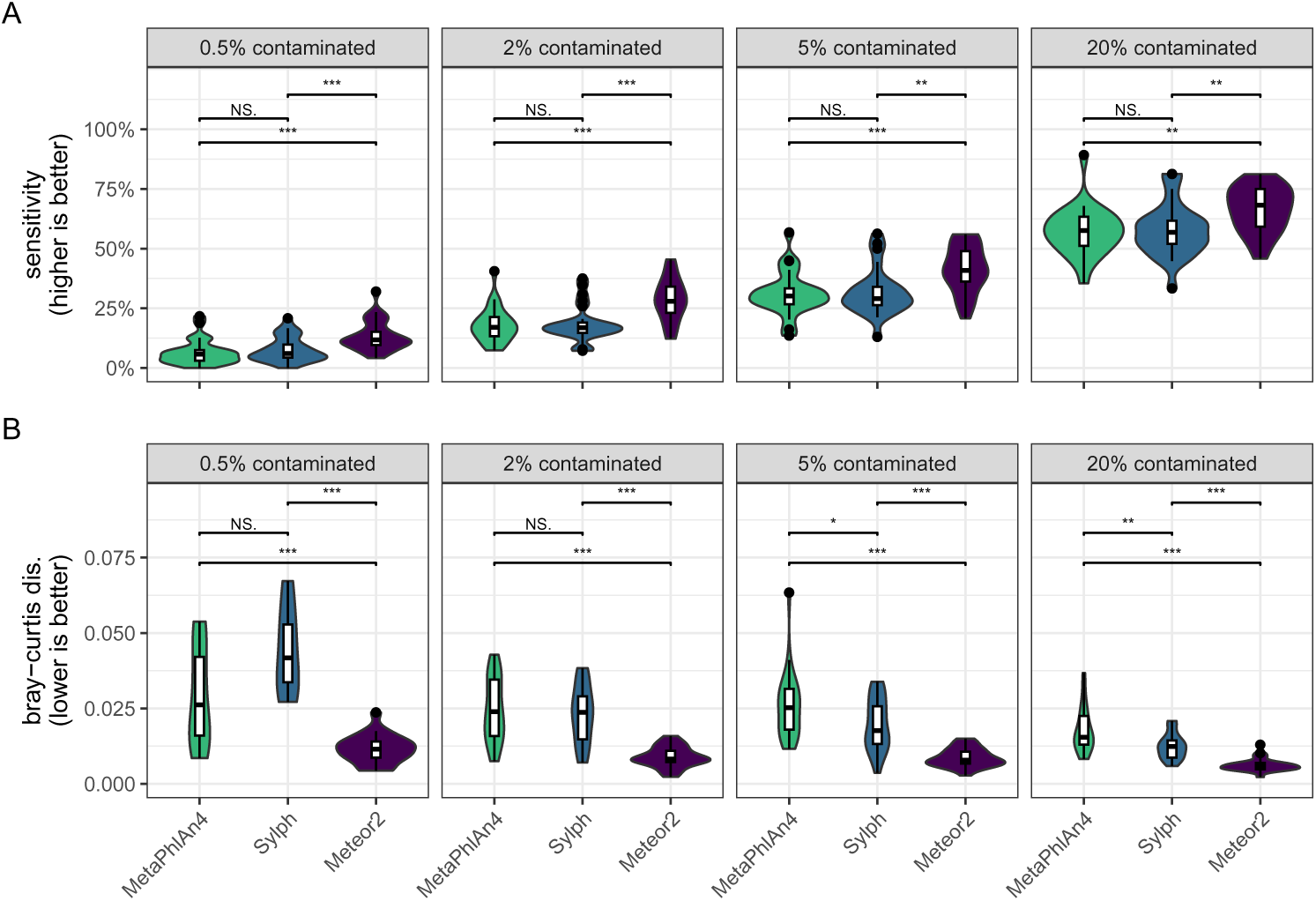
Performance of taxonomic profilers for detecting and quantifying species introduced by contamination. The 25 semi-simulated sample pairs (5M paired-end reads) with varying contamination rates (0.5%, 2%, 5%, 20%) were analyzed using MetaPhlAn4 (green), sylph (blue), and Meteor2 (purple) to generate species abundance profiles. (A) Sensitivity of each profiler in detecting species specifically introduced by contamination. Meteor2 consistently achieves higher sensitivity across all contamination rates. (B) Bray-Curtis dissimilarity between observed and theoretical abundances for contaminant species detected by each profiler. Overall, Meteor2 demonstrates superior performance for the quantification of contaminant species. Statistical comparisons were made using Wilcoxon signed-rank tests, with significance denoted as follows: ***p <0.001, **p <0.01, *p <0.05, NS (not significant).

**Fig. S5.**
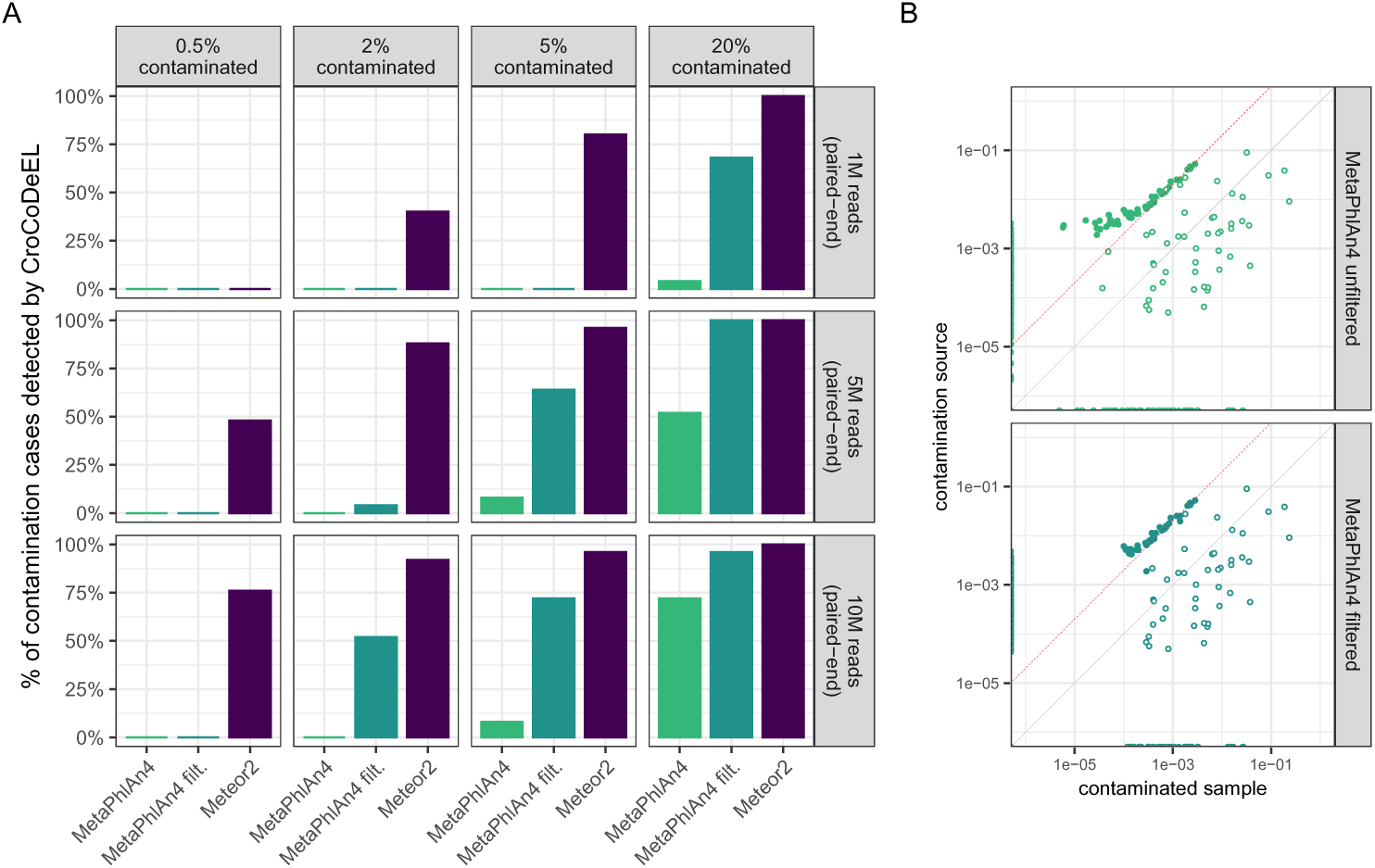
Filtering low abundance species in MetaPhlAn4 profiles improves CroCoDeEL’s sensitivity. (A) CroCoDeEL’s sensitivity was evaluated using 25 semi-simulated sample pairs (see Fig. 4), considering varying contamination rates (0.5%, 2%, 5%, 20%), sequencing depths (1M, 5M, 10M paired-end reads), and three types of taxonomic profiles: MetaPhlAn4 (light green), MetaPhlAn4 after filtering (dark green), and Meteor2 (purple). Filtering in MetaPhlAn4 profiles was performed by setting to zero the abundance of species that were up to 20 times more abundant than the least abundant species in each sample. (B) Log-scale scatter plots comparing species abundance profiles for a specific contamination case (2% contamination, 10M paired-end reads) using unfiltered MetaPhlAn4 output (light green, top) and MetaPhlAn4 after filtering low-abundance species (dark green, bottom). Filled points represent species introduced specifically by contamination. Theoretical contamination is indicated by the red line.

**Fig. S6.**
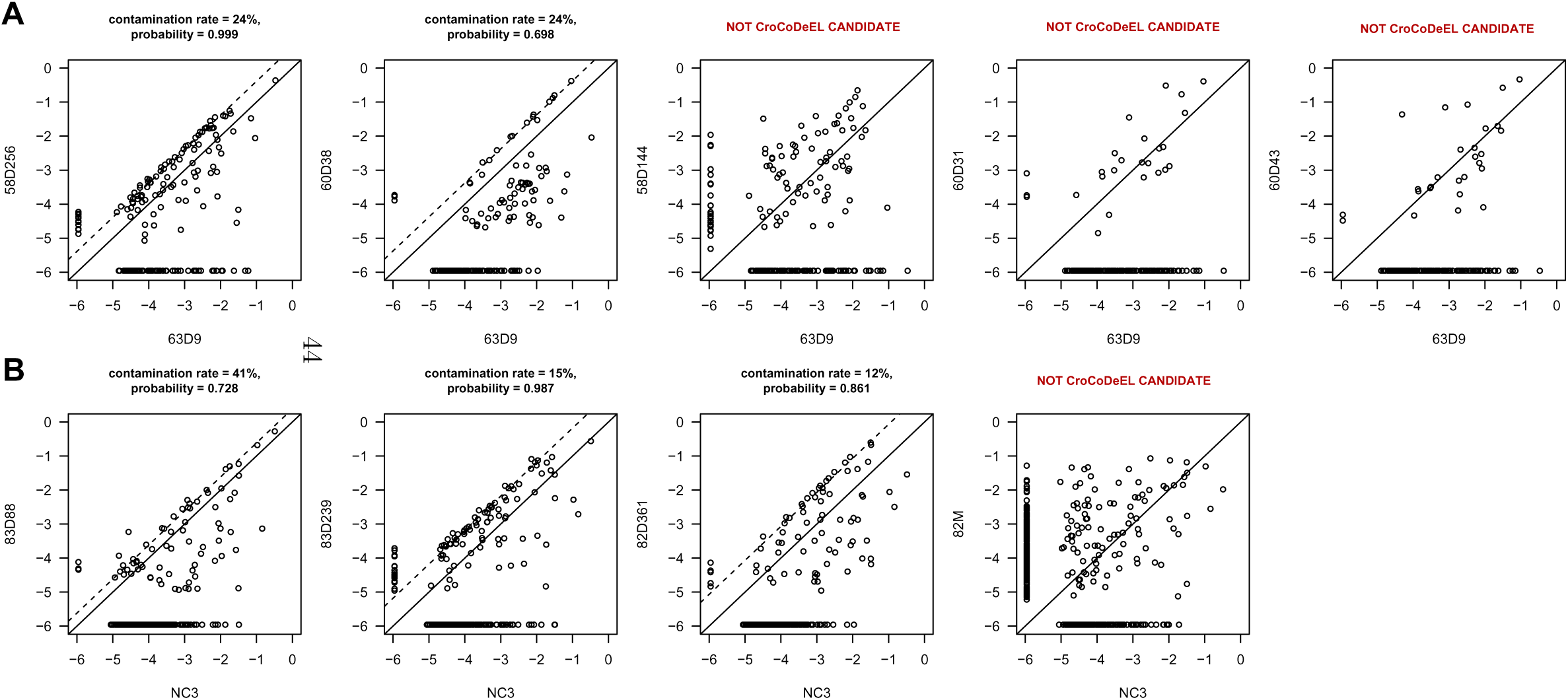
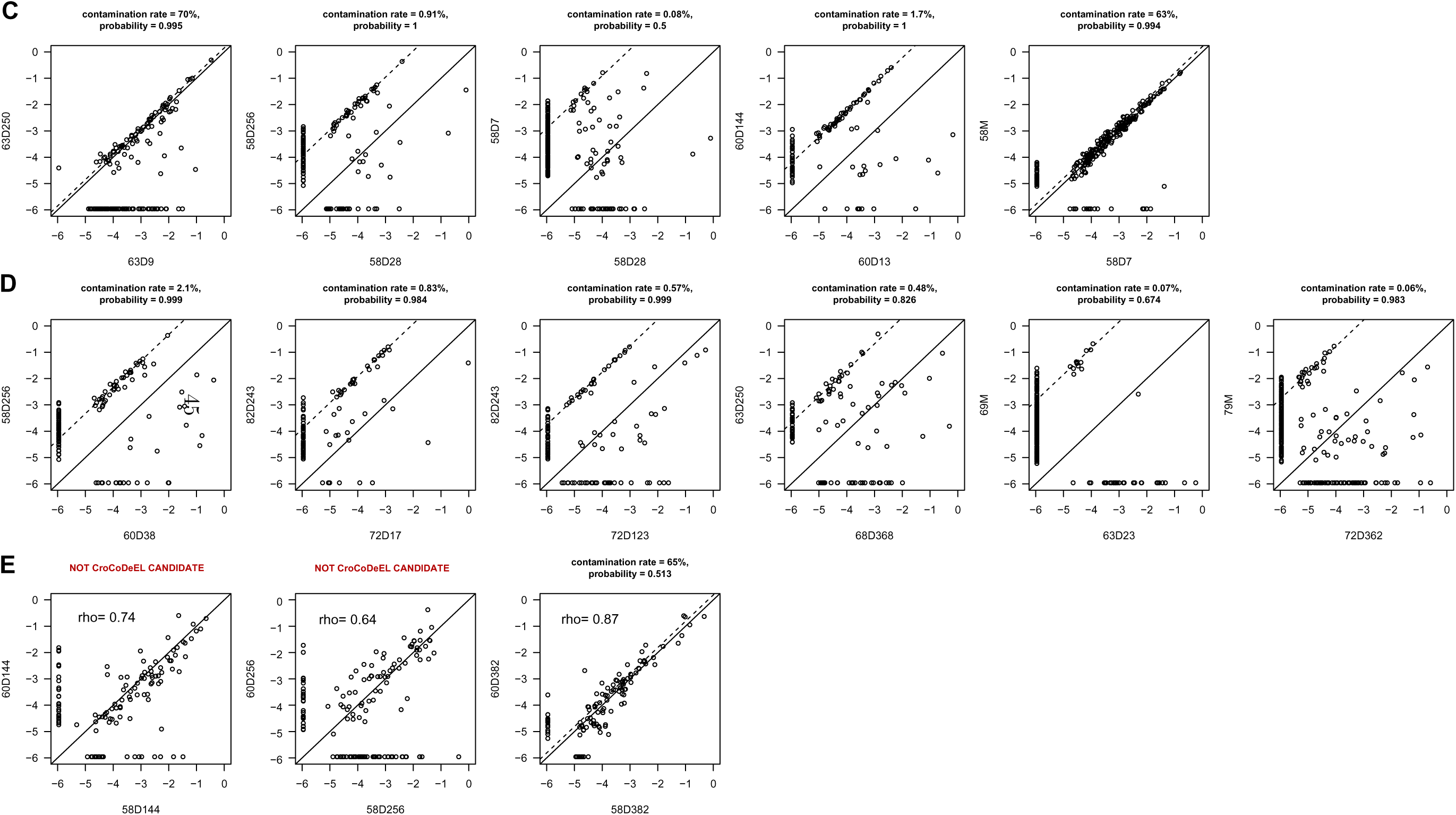
CroCoDeEL analysis of samples from plate P3 achieves greater accuracy than the strain-sharing method. (A-B) Comparison of species abundance profiles between contaminated samples (x-axis) and potential contamination sources identified by the strain-sharing method (y-axis). CroCoDeEL resolves ambiguities by accurately identifying true contamination sources and ruling out other candidates. (A) Species abundance profiles of contaminated sample 63D9 are compared with the two true contamination sources (58D256 and 60D38) identified by CroCoDeEL, along with the profiles of three other candidates identified by the strain-sharing method (physically closest on plate P3 or sharing the most strains), which CroCoDeEL exonerates. (B) Species abundance profiles of contaminated sample NC3 are compared with the three contamination sources identified by CroCoDeEL. Sample 82M is ruled out as a contamination source by CroCoDeEL. (C-D) CroCoDeEL identifies contamination events that were previously overlooked. (C) Species abundance profiles of contamination events involving related samples, detected only by CroCoDeEL, which could not be analyzed with the strain-sharing method due to natural strain sharing. (D) Species abundance profiles of contamination events involving unrelated samples, detected only by CroCoDeEL but missed by the strain-sharing method, likely due to insufficient sensitivity. (E) Comparison of species abundance profiles between twin infants 58 and 60 at different time points. CroCoDeEL reports a contamination event for sample 58 at day 382. Although the species abundance profiles are similar (as indicated by the Spearman correlation coefficient), no clear contamination line is observed. A similarity in the microbiota of these twins is observed at earlier time points, in particular at D144, but it is much less pronounced.

**Fig. S7.**
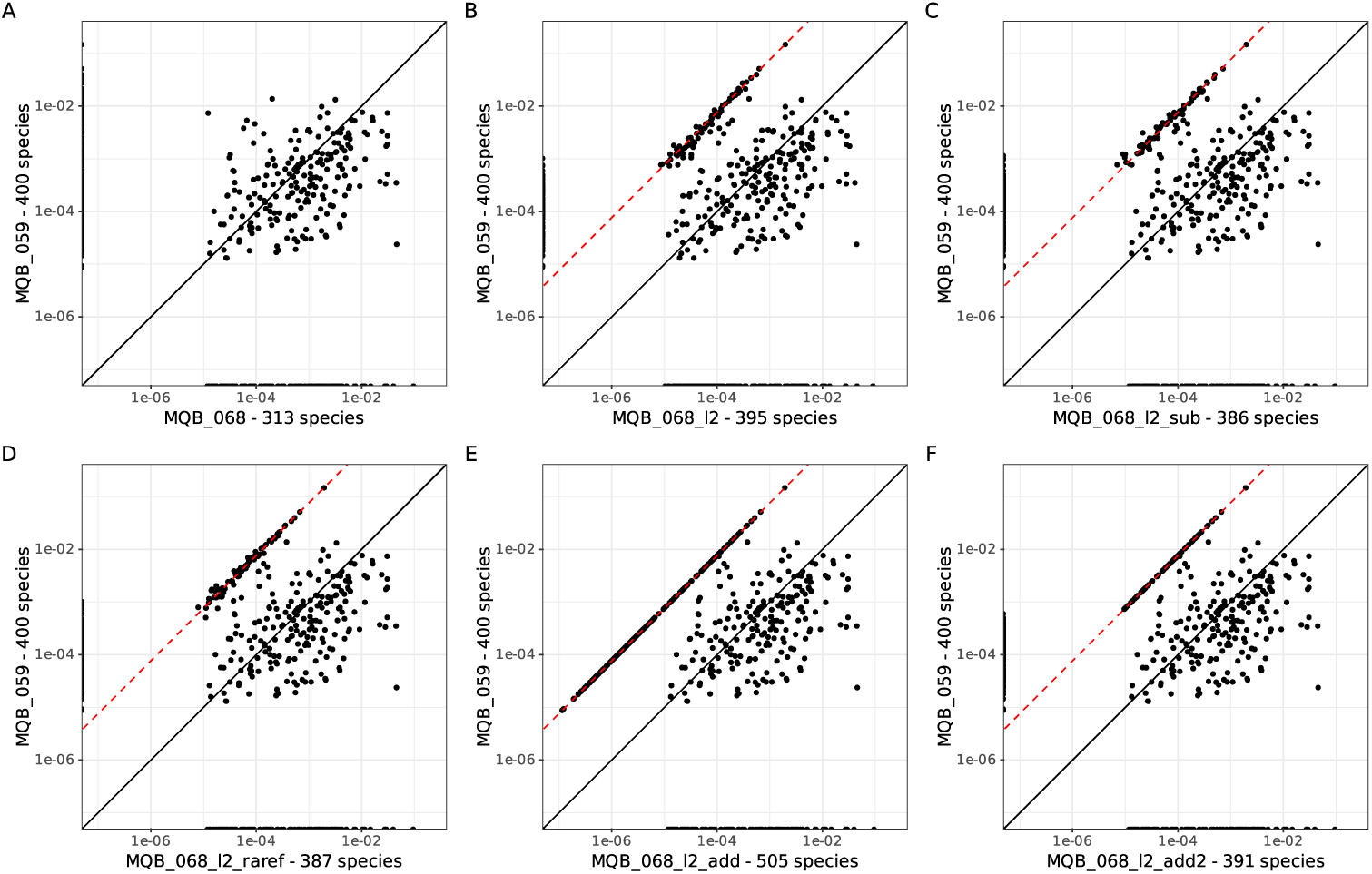
Comparison of the species abundance profiles in real and simulated contaminated samples. Scatter plots illustrate the species abundance profiles of sample pairs on a logarithmic scale. The red dashed line, when applicable, represents the contamination line. The x- and y-axis titles indicate the species richness of each sample. (A) Abundance profiles of MQB_059 and MQB_068, two real uncontaminated human fecal metagenomes. (B) Abundance profiles of MQB_059 and MQB_068_l2, a real contaminated sample created by deliberately mixing MQB_059 and MQB_068 DNA at a 99:1 ratio. The estimated contamination rate for MQB_068_l2 was 1.31%. (C) Abundance profiles of MQB_059 and MQB_068_l2_sub, a simulated contaminated sample generated by mixing sequencing reads from MQB_068 and MQB_059. (D) Abundance profiles of MQB_059 and MQB_068_l2_raref, a simulated contaminated sample created by rarefying gene count profiles from MQB_068 and MQB_059. (E) Abundance profiles of MQB_059 and MQB_068_l2_add, a simulated contaminated sample obtained by linearly combining species abundance profiles from MQB_068 and MQB_059. (F) Abundance profiles of MQB_059 and MQB_068_l2_add2, a simulated contaminated sample created by linearly combining species abundance profiles from MQB_068 and MQB_059, followed by filtering out subdominant contaminant species assumed to be undetectable. Simulation approaches based on mixing sequencing reads (C) or rarefying gene counts (D) produced species abundance profiles similar to the real contaminated sample. However, linear combinations of species abundance profiles (E) resulted in unrealistically high species richness, suggesting excessive sensitivity to subdominant species. Filtering out subdominant species (F) corrected species richness but led to overly precise estimation species abundance, shown by the very low dispersion of the contamination line.

